# Hallmarks of *Alpha*- and *Betacoronavirus* non-structural protein 7+8 complexes

**DOI:** 10.1101/2020.09.30.320762

**Authors:** Boris Krichel, Ganesh Bylapudi, Christina Schmidt, Clement Blanchet, Robin Schubert, Lea Brings, Martin Koehler, Renato Zenobi, Dmitri Svergun, Kristina Lorenzen, Ramakanth Madhugiri, John Ziebuhr, Charlotte Uetrecht

## Abstract

Coronaviruses infect many different species including humans. The last two decades have seen three zoonotic coronaviruses with SARS-CoV-2 causing a pandemic in 2020. Coronaviral non-structural proteins (nsp) built up the replication-transcription complex (RTC). Nsp7 and nsp8 interact with and regulate the RNA-dependent RNA-polymerase and other enzymes in the RTC. However, the structural plasticity of nsp7+8 complex has been under debate. Here, we present the framework of nsp7+8 complex stoichiometry and topology based on a native mass spectrometry and complementary biophysical techniques of nsp7+8 complexes from seven coronaviruses in the genera *Alpha*- and *Betacoronavirus* including SARS-CoV-2. Their complexes cluster into three groups, which systematically form either heterotrimers or heterotetramers or both, exhibiting distinct topologies. Moreover, even at high protein concentrations mainly heterotetramers are observed for SARS-CoV-2 nsp7+8. From these results, the different assembly paths can be pinpointed to specific residues and an assembly model is proposed.

## Introduction

Seven coronaviruses (CoV) from six coronavirus species are known to cause infections in humans. While four of these viruses (HCoV-229E, HCoV-NL63, HCoV-OC43, HCoV-HKU1) predominantly cause seasonal outbreaks of (upper) respiratory tract infections with mild disease symptoms in most cases, three other coronaviruses (SARS-CoV, MERS-CoV and SARS-CoV-2) of recent zoonotic origin are associated with lower respiratory tract disease including acute respiratory distress syndrome (ARDS) [1–3]. SARS-CoV-2 is the etiologic agent of COVID-19, a respiratory disease with a wide spectrum of clinical presentations and outcomes. First detected in December 2019, it quickly became pandemic with numbers still growing (>30 million confirmed cases, >1,000,000 deaths, by end September 2020) [4–6]. COVID-19 caused major perturbations of historical dimensions in politics, economics and healthcare. Moreover, coronaviruses are important, widespread animal pathogens as illustrated by feline intestine peritonitis virus (FIPV) causing a severe and often fatal disease in cats [7] or porcine coronaviruses [8], such as transmissible-gastroenteritis virus (TGEV) or porcine epidemic diarrhea virus (PEDV), the latter causing massive outbreaks and economic losses in swine industry.

The viral replication machinery is largely conserved across the different coronavirus species from the four currently recognized genera *Alpha*-, *Beta*-, *Gamma*- and *Deltacoronavirus* (subfamily *Orthocoronavirinae*, family *Coronaviridae*) [9]. The key components are generally referred to as nonstructural proteins (nsp) and encoded by the viral replicase genes (ORFs 1a and 1b) and translated as parts of the replicase polyproteins pp1a (nsp1-11) or pp1ab (nsp1-16). Translation of the ORF1b-encoded *C*-terminal part of pp1ab requires a ribosomal (−1)-frameshift immediately upstream of the ORF1a stop codon. Two proteases called PLpro (one or two protease domains in nsp3) and Mpro (also called 3CLpro or nsp5) facilitate polyprotein processing into 16 (sometimes 15) mature nsps. The majority of these nsps form a membrane-anchored, highly dynamic protein-RNA machinery, the replication-transcription complex (RTC), which mediates replication of the ~30 kb single-strand (+)-sense RNA genome and production of subgenomic mRNAs [9, 10].

The main CoV-RTC building block is the fastest known RNA-dependent RNA-polymerase (RdRp) residing in the nsp12 C-terminal domain [11]. For RdRp activity, nsp12 requires binding to its cofactors nsp7 and nsp8 [12]. Recently, high-resolution structures illuminated two binding sites at nsp12, the first for an nsp7+8 (1:1) heterodimer and the second for a single nsp8 [13–16]. For *in vitro* RdRp activity assays, different methods were used to assemble the polymerase complex [11, 17, 18]. So far, the highest processivity *in vitro* was obtained by mixing nsp12 with a flexibly linked nsp7L8 fusion protein.

Recently, we reported that SARS-CoV nsp7 and nsp8 form a heterotetramer (2:2) in solution, in which nsp7 subunits have no self-interaction and rather sandwich an nsp8 scaffold with putative head-to-tail interactions [19]. Current knowledge of full-length nsp7+8 complexes is mainly based on two X-ray crystal structures, each of which displays a different quaternary conformation. First, a SARS-CoV nsp7+8 (8:8) hexadecamer is assembled from four (2:2) heterotetramers with similar topologies but two distinct conformations, T1 and T2, which are both consistent with our in solution structure [19] [20]. Second, in a feline coronavirus (FIPV) nsp7+8 (2:1) heterotrimer, nsp8 is associated to two nsp7 molecules that self-interact [21]. Moreover, structures of SARS-CoV and SARS-CoV-2 with *N*-terminally truncated forms of nsp8, thus lacking the self-interaction domain, revealed heterotetrameric nsp7+8 complexes around an nsp7 scaffold [22, 23].

Current knowledge of coronavirus nsp7+8 complexes suggests a remarkable architectural plasticity but is unsupportive of deducing common principles of complex formation. Moreover, it is unknown if the quaternary structure of nsp7+8 is conserved within a given coronavirus species or between genera. To fill these knowledge gaps, we analyzed nsp7+8 complexes derived from seven viruses of the *Alpha*- and *Betacoronavirus* genera, including a range of human coronaviruses, namely SARS-CoV, SARS-CoV-2, MERS-CoV and HCoV-229E (Table S 2). We used native mass spectrometry (MS) to illustrate the landscape of nsp7+8 complexes *in vacuo*, collision induced dissociation tandem MS (CID-MS/MS) to reconstruct complex topology and complementary methods to verify the results [24, 25]. Our findings reveal distinct sets of nsp7+8 complexes for the different CoV species. The results hint at the properties that lead to complex heterogeneity and suggest common principles of complex formation based on two conserved binding sites.

## Experimental Results

### Native MS illustrates the landscape of nsp7+8 complexes

To ensure authentic nsp7 and nsp8 N- and C-termini, which allow for optimal nsp7+nsp8 complex assembly, the proteins are expressed as nsp7-8-His_6_ polyprotein precursors and cleaved by their cognate protease M^pro^ (Figure S 1). Native MS provides an overview of mass species in solution, while CID-MS/MS confirms the stoichiometry of protein complexes. Distinct oligomerization patterns of nsp7+8 (1:1) heterodimers, (2:1) heterotrimers and (2:2) heterotetramers in the different CoV allowed us to categorize their nsp7+8 complexes into three groups (Figure 1, Figure S 2). SARS-CoV and SARS-CoV-2 (species *Severe acute respiratory syndrome-related coronavirus*, genus *Betacoronavirus*) represent nsp7+8 group A complex formation pattern (Figure 1 A). Consistent with our previous work, SARS-CoV nsp7+8 complexes exist primarily as a heterotetramer comprising two copies of each nsp7 and nsp8 (2:2) [19]. Expectedly, SARS-CoV-2 nsp7+8 form identical (2:2) complexes given the high sequence identity of 97.5 % in the nsp7-8 region (Table S 2). Next, relative peak intensities in native MS of nsp7+8 complexes are converted in a semi-quantitative analysis into abundances of complex species [26]. The heterodimer (2-4 %) is much less abundant than the heterotetramer (96-98 %) suggesting high affinity and hence efficient conversion of heterodimeric intermediates into heterotetramers. Hence, group A only forms two types of nsp7+8 complexes, heterodimers (1:1) and -tetramers (2:2), with the latter clearly being predominant.

**Figure 1:**
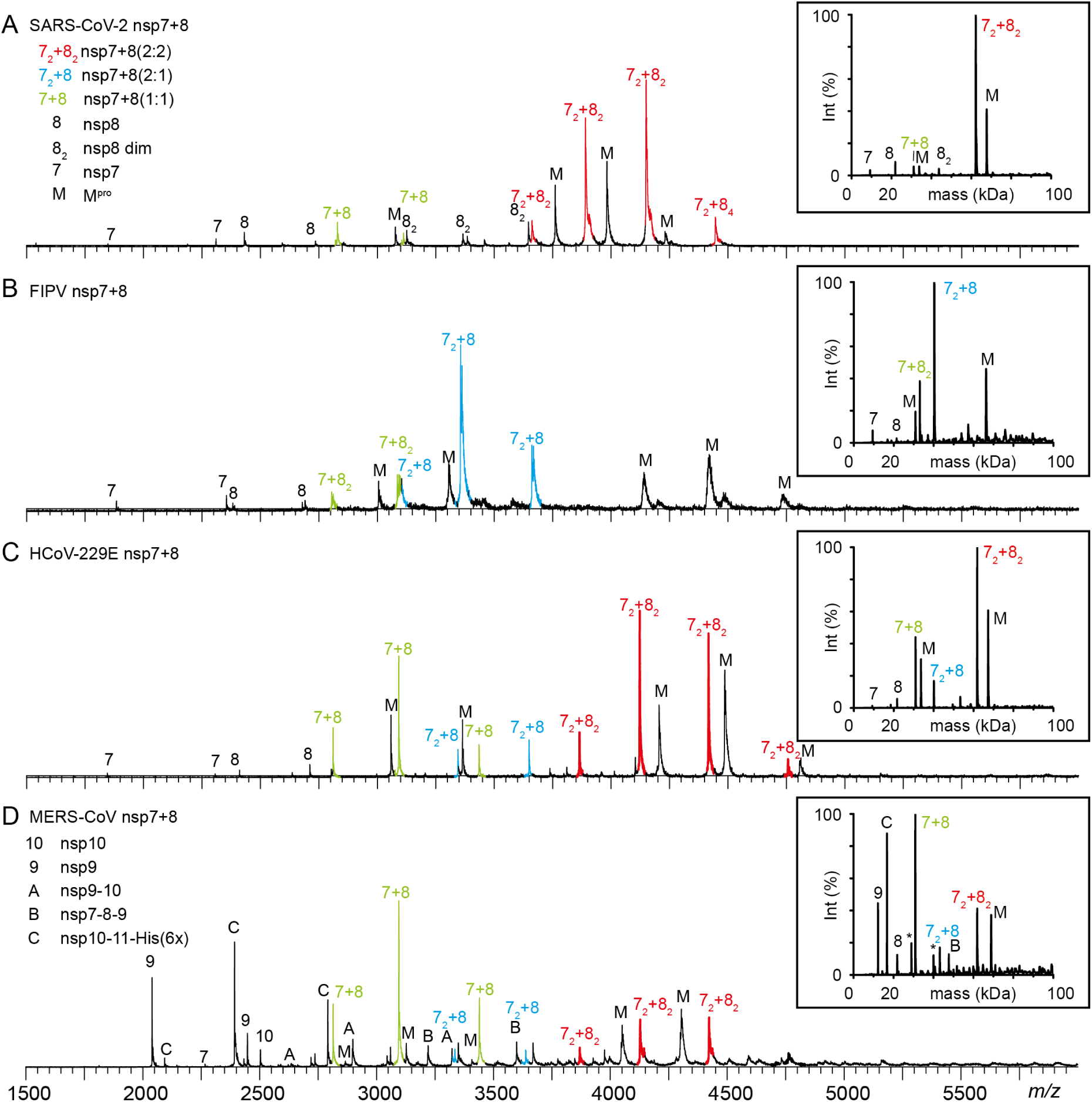
Three complexation groups formed by CoV nsp7+8. Representative mass spectra showing distinct nsp7+8 complexation patterns that were classified into the three groups A, B and AB. Complex formation triggered by M^pro^ (M) mediated cleavage of 15 μM CoV nsp7-8-His_6_ or MERS-CoV nsp7-11-His_6_ precursors in 300 mM ammonium acetate (AmAc), 1 mM DTT, pH 8.0. (**A**) SARS-CoV-2 representing group A forms heterotetramers (nsp7+8 (2:2), red), (**B**) FIPV from group B forms heterotrimers (nsp7+8 (2:1), blue) and (**C**) HCoV-229E from group AB forms both complex stoichiometries (2:2 and 2:1). (**D**) In case of MERS-CoV, also group AB, cleavage of the longer nsp7-11-His_6_ precursor resulted in additional processing intermediates (labeled A, B, C). In all three complexation groups, nsp7+8 (1:1) heterodimeric intermediates are observed (green). For spectra of all seven CoV nsp7+8 complexes see Figure S 2.

In FIPV and TGEV from the species *Alphacoronavirus 1*, genus *Alphacoronavirus*, nsp7 and nsp8 proteins share a sequence identity of 93.9% (Table S 2). Their nsp7+8 complexes are assigned to group B forming predominantly nsp7+8 (2:1) heterotrimers (83 %) and to a lesser extent heterodimers (1:1) (~17 %) (Figure 1 B). An nsp7+8 (2:1) heterotrimeric structure has previously been reported for FIPV but not for TGEV or any other CoV. The association of a single nsp8 with two nsp7 indicates that group B nsp7+8 complexes lack the ability to form tetramers around an nsp8 scaffold.

The third oligomerization pattern is observed for nsp7+8 of HCoV-229E and PEDV, which represent different species in the genus *Alphacoronavirus*. They share only 70.9 % sequence identity in the nsp7-8 region and even less (42-62 %) with the other CoV species examined (Table S 2). PEDV and HCoV-229E nsp7+8 form three major types of oligomers with slightly different efficiencies: heterodimers (1:1), heterotrimers (2:1) and heterotetramers (2:2) (HCoV-229E: 20 %, 12 % and 69 %; PEDV: 52 %, 6 % and 42 %, respectively) (Figure 1 C). By forming both, heterotrimers and -tetramers, these complexes combine properties described above for groups A and B, and are hence categorized into a separate group named accordingly AB. This begs the question whether assembly pathways and structures of heterotetramers in group A and AB are similar. Either, two heterodimers form a heterotetramer around an nsp8 scaffold as in group A [19] or alternatively the heterotrimer recruits another nsp8 subunit to the complex, thus employing an nsp7 core [21]. The latter pathway has recently been reported for SARS-CoV-2 nsp7+8 heterotetramers containing *N*-terminally truncated nsp8 [23].

Additionally, nsp7+8 complexation after M^pro^ mediated cleavage of a MERS-CoV nsp7-11-His_6_ precursor is compared (Figure 1 D). This larger precursor, comprising nsp7, nsp8, nsp9 nsp10 and nsp11, behaves similar to nsp7-9-His_6_ (Figure S1) and is used because initial attempts to cleave nsp7-8 only constructs failed. Proteolytic processing of this polyprotein precursor leads to cleavage intermediates (Figure 1 D). Such processing intermediates have been proposed to occur intracellularly and to function distinctly from the individual nsps in e.g. regulation of RTC assembly and viral RNA synthesis [27]. Here, signal intensities of these intermediates provide insights into the processing sequence. Surprisingly, the dominant intermediate is nsp10-11-His_6_, despite the small size of nsp11 and a hence expected high accessibility of the nsp10/11 cleavage site. Therefore, slow cleavage and prolonged presence of an nsp10-11 intermediate may have functional implications warranting further studies. Notably, in many CoV polyproteins the nsp10/11 and/or nsp10/12 cleavage sites contain replacements (Pro in MERS-CoV) of the canonical P2 Leu residue conserved throughout most M^pro^ cleavage sites, suggesting that slow or incomplete cleavage is beneficial for these particular sites. Moreover, this cleavage site has different C-terminal contexts in the two CoV replicase polyproteins, nsp10-11 in pp1a and nsp10-12 in pp1ab. While the structure of the small nsp11 (~1.5 kDa) is unknown, nsp12 is a large folded protein (~105 kDa), which potentially improves the accessibility of the nsp10/12 site for M^pro^. Similar effects have been observed for the nsp8/9 cleavage site, which is efficiently cleaved in the protein but not in peptide substrates [19, 28]. The question remains if unprocessed nsp10-11 and/or nsp10-12 intermediates exist in virus-infected cells for prolonged times to fulfill specific functions. Other detected intermediates are nsp7-8-9 and nsp9-10 lacking nsp11-His_6_. Particularly, the nsp9-10 intermediate has not been identified in our analysis of SARS-CoV nsp7-10 processing, suggesting differences in the *in vitro* processing order between SARS-CoV and MERS-CoV.

MERS-CoV nsp7+8 forms heterodimers (1:1), -trimers (2:1) and -tetramers (2:2) (73 %, 8 % and 19 %, respectively), thus demonstrating a group AB complexation pattern. However, we cannot confirm the heterotrimer (2:1) formation by CID-MS/MS due to spectral complexity.

Moreover, because of incomplete cleavage as is evident from the cleavage intermediates, signals assigned to the nsp7+8 heterodimer likely overlap with signals of unprocessed nsp7-8. Thus, complete cleavage of nsp7-8 could shift the peak fractions from heterodimer to heterotrimer or -tetramer.

### Homodimerization of subunits and precursors

In the mass spectra of nsp7+8 complexes, monomers and homodimers of nsp7 and nsp8 are also observed. While nsp7 homodimers are identified for all seven CoV species tested, nsp8 homodimers are only detected for SARS-CoV and SARS-CoV-2, which belong to group B forming exclusively nsp7+8 heterotetramers around a dimeric nsp8 scaffold (Figure S 3). Moreover, the oligomeric states of the different uncleaved nsp7-8 precursors are probed. Notably, precursors from group B CoVs are mostly monomeric, whereas precursors from group AB and A CoVs are in varying equilibria between monomers and dimers (Figure S 4). The different oligomerization propensities of precursors suggest that molecular interactions driving dimerization of nsp7-8 precursors could critically affect subsequent nsp7+8 oligomerization. For SARS-CoV and SARS-CoV-2, the nsp7-8 dimer affinity is low as two-fold dilution to 9 μM shifted the equilibrium towards a monomeric state (Figure S 5). This is in line with our previous findings [19], in which *C*-terminally extended SARS-CoV nsp7-9-His_6_ and His_6_-nsp7-10 polyprotein constructs were mainly monomeric, suggesting that the presence of the extra *C*-terminal sequence further destabilizes an already weak dimerization.

### Collision-induced dissociation reveals complex topology

To deduce the complex topology in the different groups of nsp7+8 interaction patterns, we applied CID-MS/MS using successive subunit dissociations to dissect conserved interactions. CID-MS/MS of the HCoV-229E nsp7+8 heterotetramer (2:2) reveals two dissociation pathways, in which first one nsp7 subunit is ejected from the complex followed by another nsp7 or an nsp8 subunit. After two consecutive losses, the product ions are nsp7+8 (1:1) and nsp8_2_ dimers, providing evidence for specific subunit interfaces in the complex (Figure 2 A). From these results, the complex topology is deduced as a heterotetramer based on an nsp8_2_ dimer scaffold, in which each nsp8 binds only one nsp7 subunit. Strikingly, this is identical to our previously reported SARS-CoV nsp7+8 heterotetramer (2:2) architecture [19]. In fact, all nsp7+8 (2:2) heterotetramers of groups A and AB (SARS-CoV-2, SARS-CoV, PEDV, HCoV-229E and MERS-CoV) resulted in similar dissociation pathways, subunit interfaces and topology maps, suggesting that these structures are similar across these diverse CoVs.

Next, the dissociation pathway of the HCoV-229E nsp7+8 (2:1) heterotrimers is monitored in CID-MS/MS (Figure 2 B). After ejection of one nsp7 or nsp8 subunit, product dimers of nsp7+8 (1:1) and nsp7_2_ are detected, indicating specific subunit interfaces between nsp7:nsp8 and nsp7:nsp7. Again, similar dissociation pathways and subunit interfaces are found for group B and AB heterotrimers (FIPV, TGEV, HCoV-229E and PEDV). Topological reconstructions reveal a heterotrimer forming a tripartite interaction between one nsp8 and two nsp7 subunits. These results agree with the reported X-ray structure of FIPV nsp7+8 [21] and indicate that heterotrimers of these CoVs species have similar arrangements. In turn, this implies that heterotrimers and heterotetramers follow distinct assembly paths.

**Figure 2:**
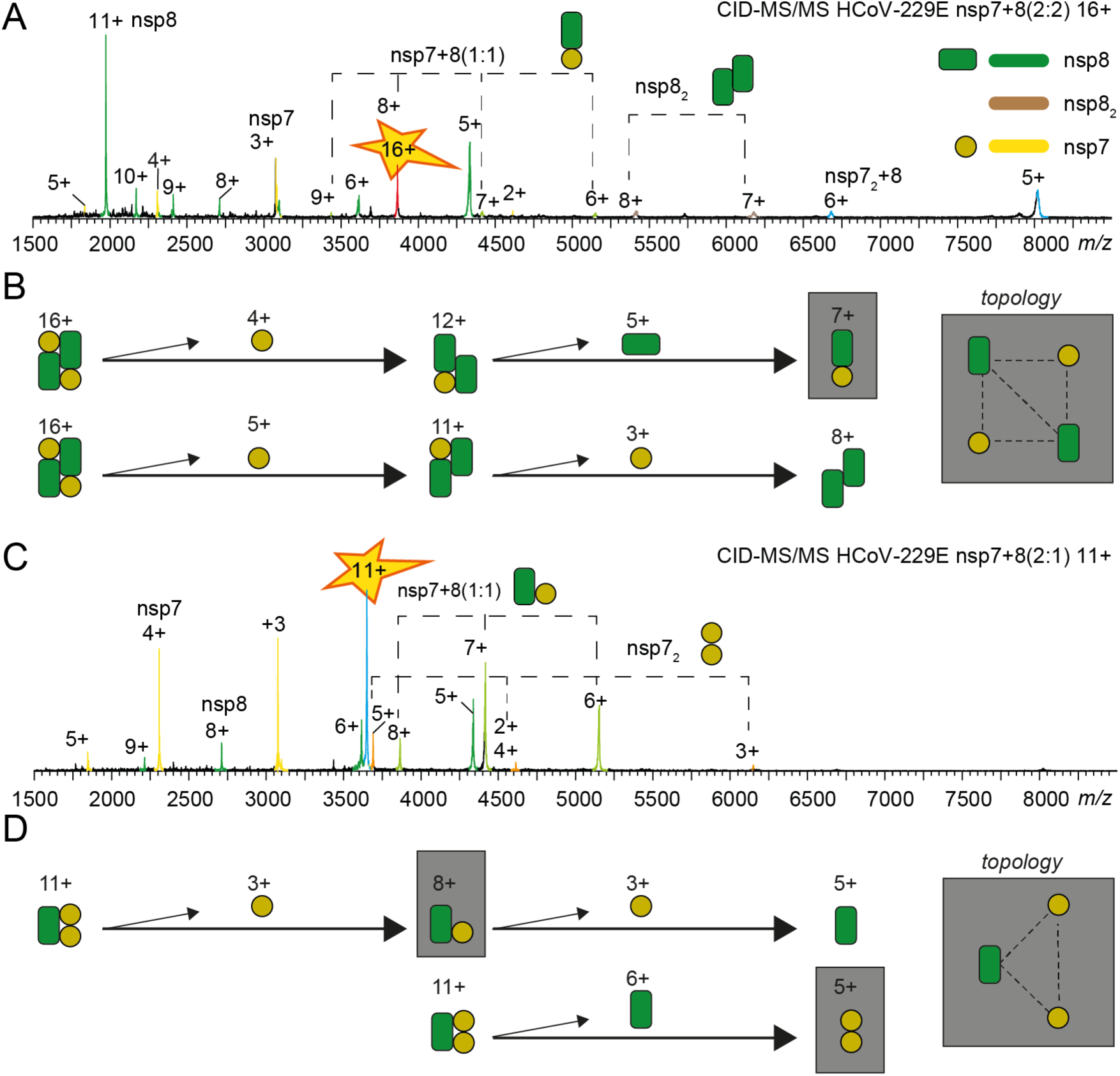
Gas-phase dissociation reveals complex topology. CID-MS/MS product ion spectra (**A** and **C**), dissociation pathways and topology maps (**B** and **D**) for HCoV-229E nsp7+8 (2:1) heterotrimers and (2:2) heterotetramers are shown. With increasing collisional voltage protein complexes are successively stripped from their subunits revealing alternative dissociation pathways. The remaining dimeric species expose direct subunit interactions in the nsp7+8 complexes (grey boxes). Charge states are labeled. (**A** and **B**) The heterotetramers (2:2) undergo two consecutive losses resulting in dimeric product ions of nsp7+8 (1:1) and nsp8_2_. These products indicate that nsp7:nsp8 and nsp8:nsp8 have direct interfaces in heterotetramers. (**C** and **D**) HCoV-229E heterotrimers dissociate into the dimeric products nsp7+8 (1:1) and nsp7_2_ indicating direct interfaces between nsp7:nsp8 and nsp7:nsp7 in heterotrimers. All CoV heterotrimers follow similar dissociation pathways, also all CoV heterotetramers follow a common dissociation route, which allow a topological reconstruction of two distinct complex architectures (Figure S 6–Figure S 10).

### Chemical cross-linking confirms the formation of specific complexes

To further support the native MS results, which relies on spraying from volatile salt solutions (e.g. ammonium acetate, AmAc), complementary methods compatible with conventional buffers supplemented with sodium chloride are applied. To provide additional evidence for specific nsp7+8 complex formation, the FIPV and HCoV-229E nsp7+8 complexes are stabilized via cross-linking with glutaraldehyde and subjected to XL-MALDI MS (Cross-linking Matrix-assisted laser ionization MS) (Figure S 11). Peak areas in the MALDI mass spectra are assigned to FIPV nsp7+8 heterodimer, heterotrimer and –tetramer (8.5 %, 9.5 % and 4.2 %, respectively), and HCoV-229E nsp7+8 heterodimer, heterotrimer and –tetramer (6.7 %, 5.3 % and 6.0 %, respectively). The results suggest a higher abundance of nsp7+8 heterodimer and -trimer complexes in FIPV than in HCoV-229E, while HCoV-229E contains more heterotetramers. This largely agrees with the results from native MS. However, the MALDI mass spectra show high background of virtually all possible nsp7+8 stoichiometries (<200,000 *m*/*z*), probably due to over-cross-linking with the rather unspecific glutaraldehyde.

To refine these results, nsp7+8 complexes are stabilized with the amine specific cross-linker BS^3^ and analyzed by SDS-PAGE (Figure S 12). Multiple stoichiometries are identified with a few prominent bands highlighting the main complexes generated. These bands are assigned to SARS-CoV nsp7+8 heterodimers and -tetramers, FIPV nsp7+8 heterodimers and -trimers and HCoV-22E nsp7+8 heterodimers, heterotrimers and -tetramers of HCoV-229E, providing additional support for the classification of nsp7+8 complexes into groups A (SARS-CoV), B (FIPV) and AB (HCoV-229E).

### Light scattering provides insights into complexation at high protein concentrations

**Figure 3:**
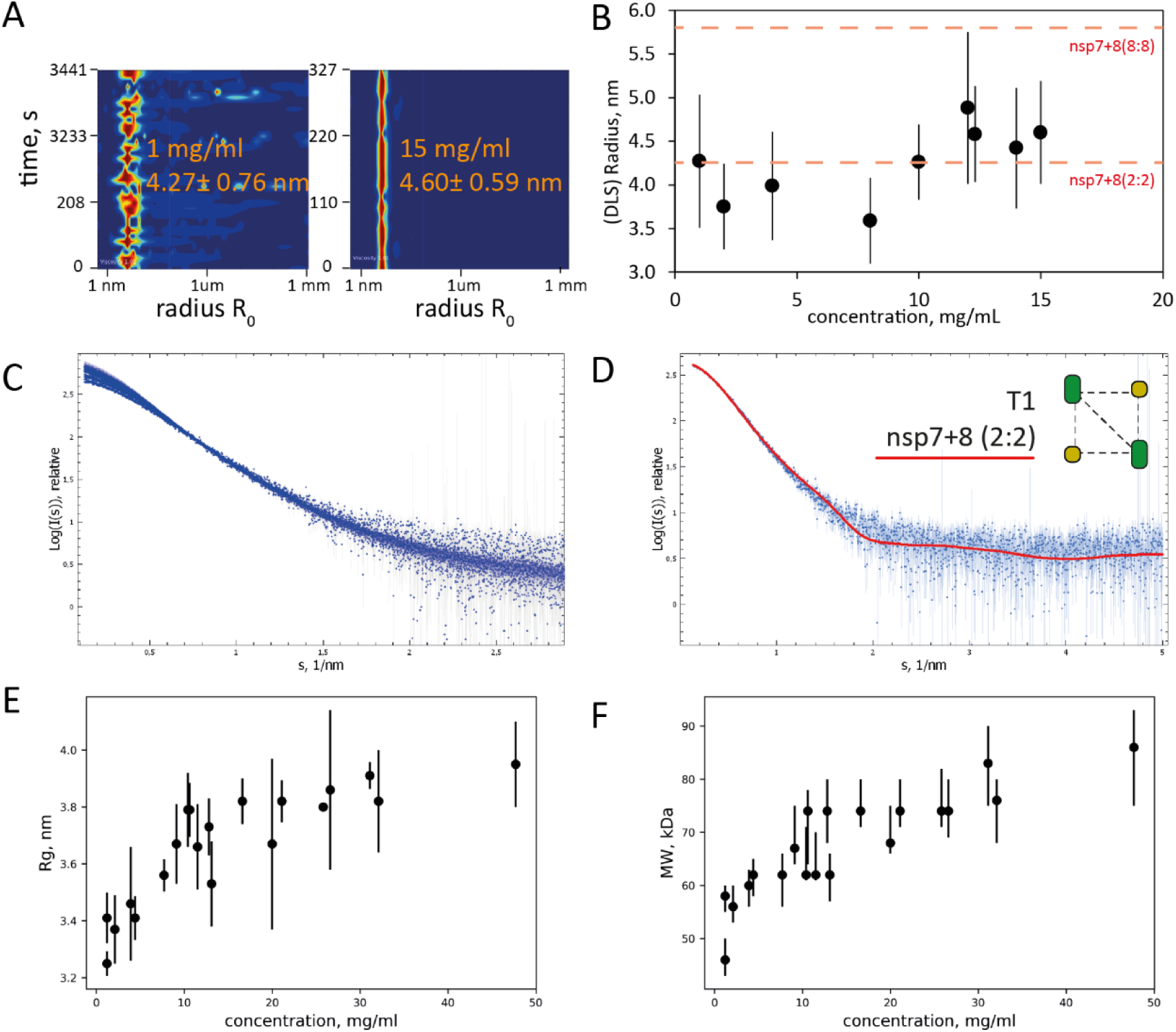
DLS and SAXS reveal oligomeric state of SARS-CoV-2 nsp7+8 at higher protein concentrations. At increasing protein concentrations, the hydrodynamic radius (*R_0_*) remains stable but becomes more homogenous in DLS. (**A**) shows two exemplary DLS plots (1 mg/ml; 15 mg/ml) and (**B**) how the radius *R_0_* develops with increasing protein concentration. Theoretical hydrodynamic radii (*R*_0,theo_) of heterotetramer and hexadecamer candidate structures are indicated (red dashed lines). (**C**) displays SAXS curves collected at different solute concentrations and (**D**) the fit of the curve computed from T1 tetramer (red line) to the SAXS data collected at 1.2 mg/mL (blue dots with error bars). (**E**) Radius of gyration (*R_g_*) and (**F**) molecular weight estimated from the SAXS data both stabilize with increasing concentration like in DLS on values that are in agreement with the *R*_0,theo_ of the T1 nsp7+8 (2:2) heterotetramer.

To test the stoichiometry at higher protein concentrations in solution, dynamic light scattering (DLS) of SARS-CoV-2 nsp7+8 from 1 to 15 mg/mL is performed (Figure 4 A). No significant increase of the hydrodynamic radius (*R*_0_) occurrs with increasing concentration. At the same time, the measured radii become more stable and fluctuate less, which suggests a shift towards higher complex homogeneity and a reduced fraction of free nsp7 and nsp8.

**Figure 4:**
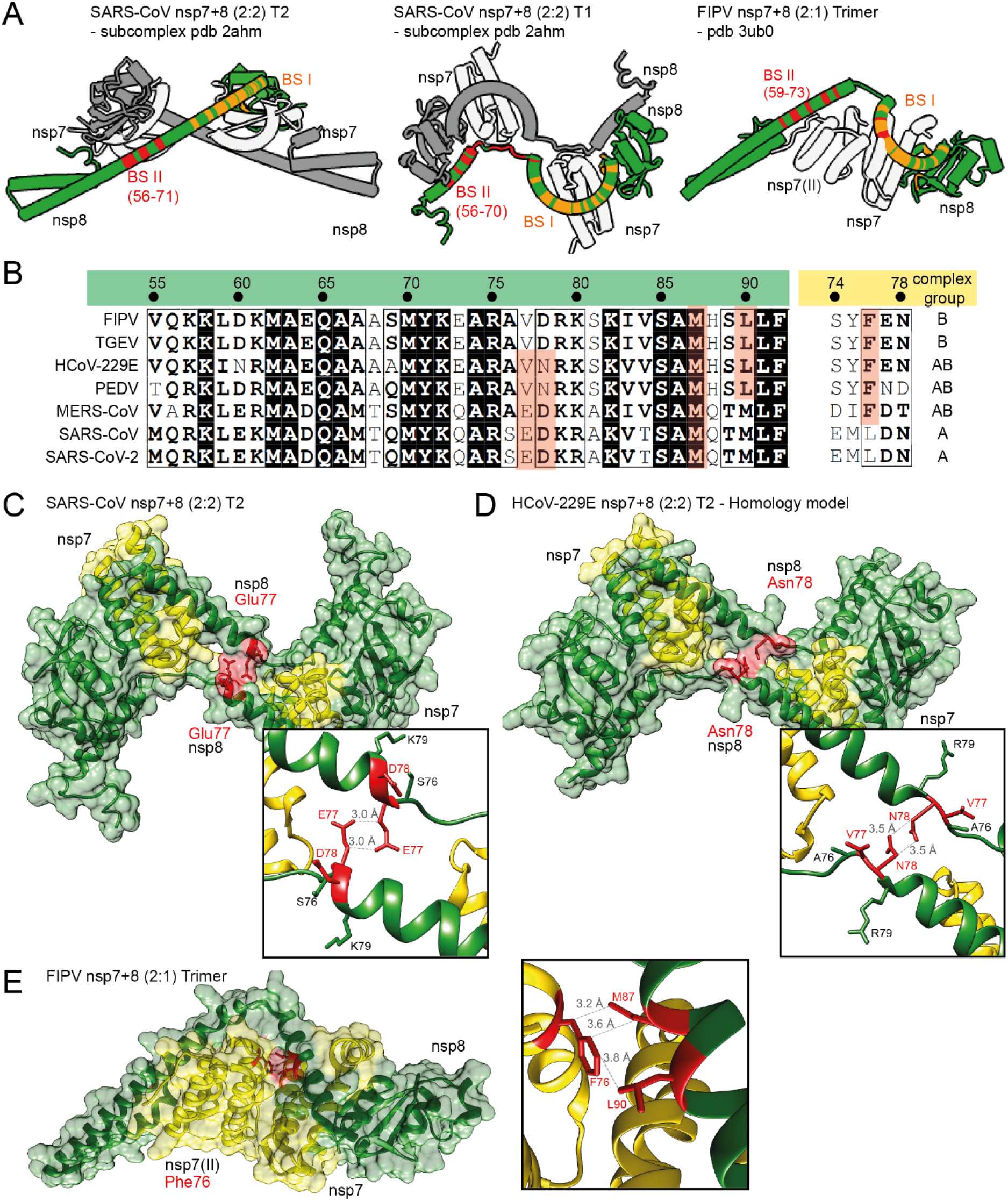
Candidate structures and sequence conservation of nsp7+8 heterotrimers and -tetramers. Candidate structures for nsp7+8 heterotetramer and –trimer are chosen based on experimental stoichiometry and topology in solution. (**A**) shows two conformers of SARS-CoV nsp7+8 (2:2) heterotetrameric subcomplexes, T1 (left) and T2 (middle), from the larger (8:8) heterohexadecamer (pdb 2ahm) and the FIPV nsp7+8 (2:1) heterotrimer (right, pdb 3ub0). Complexes exhibit two similar binding sites in nsp8, BS I (orange) and BS II (red). For simplification binding sites are only shown for one nsp8 subunit (green). BS II is additionally labelled with nsp8 residue (res) number forming the main interaction patch (BS II contact residues see Figure S 15). (**B**) Sequence alignment of nsp8 (green, res55-92) and nsp7 (yellow, res74-78) is displayed for the seven CoVs. Two heterotrimer or -tetramer specific contact sites (red) exhibit sequence conservation well in line with the complexation groups determined by native MS. (**C**) In SARS-CoV T2, nsp8 Glu77 comes into contact with nsp8 Glu77, a unique heterotetramer interaction, (**D**) which in a homology model of HCoV-229E based on T2 is replaced by nsp8 Asn78. (**E**) The FIPV heterotrimer structure shows nsp7 Phe76, binding to nsp8 Met87 and Leu90, a contact unique for the heterotrimer forming species. Insets show magnification of contact sites. Closest distances (3.0-3.8 Å) for relevant residues in contact (red) are given.

For SARS-CoV-2, no complex structure is available for full-length nsp7+8 proteins but, previously, a SARS-CoV nsp7+8 (8:8) has been reported using X-ray crystallography [20], where high protein concentrations are deployed. In order to relate the average experimental hydrodynamic radius (*R*_0,exp_ = 4.25 ± 0.61 nm) to candidate structures, the theoretical hydrodynamic radius is calculated for the SARS-CoV nsp7+8 (8:8) hexadecamer (*R*_0,theo_ = 5.80 ± 0.29 nm) and a subcomplex thereof, a putative nsp7+8 heterotetramer (2:2) in T1 conformation (*R*_0,theo_ = 4.52 ± 0.27 nm) (Figure S 13). This is the only model with full-length nsp8 that agrees with the stoichiometry and topology determined by native MS. At physiologically relevant concentrations from 1 to 10 mg/mL, the average experimental hydrodynamic radius agrees well with the theoretical hydrodynamic radius of the heterotetramer T1. Hence a heterotetramer is likely the prevailing species in solution (Figure 4 B).

To underpin the DLS results, SAXS data are collected on solutions of nsp7+8 at concentrations ranging from 1.2 to 47.7 mg/ml. The normalized SAXS intensities increase at low angles with increasing concentration (Figure 4 B and Table S 3), suggesting a change in the oligomeric equilibrium and a formation of larger oligomers. This trend is well illustrated by the evolution of the apparent radius of gyration and molecular weight of the solute determined from the SAXS data (Figure 4 E and F). The increase in the effective molecular weight, from about 50 to 80 kDa suggests that the change in oligomeric state is limited and that the tetrameric state (MW_theo_: 62 kDa) remains predominant in solution.

The SAXS data at low concentrations (<4 mg/mL) fit well the computed scattering from heterotetramer T1 but misfits appear at higher concentration (Figure 4 D, structure of T1 shown in Figure S 13 and the discrepancy χ^2^ reported in Table S 4). Mixtures of heterotetramers and hexadecamers cannot successfully fitted to the higher concentration data either (Figure S 14). To further explore the oligomeric states of nsp7+8, a dimer of T1 is used to simultaneously fit the curves collected at different concentrations by a mixture of heterotetramers and -octamers. Reasonable fits to all SAXS data are obtained with volume fractions of heterooctamers growing from 0 to 0.52 with increasing concentration (Figure S 14). Based on the flexibility of the molecule and the multiple possible binding sites between nsp7 and nsp8, it is not surprising that larger assemblies are observed at very high solute concentrations. The SAXS and DLS results provide evidence that the nsp7+8 (2:2) heterotetramer is the prevailing stoichiometry in solution at physiological concentrations (1-10 mg/mL with volume fractions between 1 and 0.7).

### Potential implications of sequence conservation on heterotrimer and -tetramer formation

To extend this analysis, we select candidate structures in agreement with the stoichiometry and topology observed (Figure S 10). For the heterotetramer, two conformers of nsp7+8 (2:2) subcomplexes, T1 and T2, of correct architecture can be extracted from the larger SARS-CoV nsp7+8 hexadecamer [20] (pdb 2ahm), (Figure S 13 A). Both conformers constitute a head to tail interaction of two nsp7+8 heterodimers mediated by an nsp8-nsp8 interface. Notably, nsp8 in T1 is more extended, revealing an almost full-length amino acid sequence (2-193), while in T2 the nsp8 *N*-terminal 35 to 55 residues are unresolved. For the heterotrimeric complexes, the only deposited structure is the FIPV nsp7+8 (2:1) heterotrimer [21] (pdb 3ub0), which agrees well with our experimental topology (Figure S 13 B).

In order to identify molecular determinants for heterotrimer or -tetramer formation, the candidate structures are examined for molecular contacts (van der Waals radius −0.4 Å). The conservation of contact residues is evaluated in a sequence alignment to identify possible determinants of different stoichiometries (Figure S 15). Notably, most amino acids lining subunit interfaces in heterodimers, -trimers and -tetramers are conserved. The interfaces in the candidate structures occupy two common structural portions of the nsp8 subunit (Figure 4 A). The first binding site (BS I) is located between the nsp8 head and shaft domain, responsible for binding of nsp7 (I) in heterodimer formation, as seen in all available high resolution structures of nsp7+8 [20–23] and the polymerase complex [13–16]. The second binding site (BS II) appears highly variable in terms of its binding partner and lies at the nsp8 elongated *N*-terminus. In fact, one largely conserved motif (res60-70) is responsible for the main contacts in the entire candidate complexes selected based on our data: nsp7+8 (2:2) T1 and T2 for the heterotetramer and nsp7+8 (2:1) for the heterotrimer. The respective sidechains take positions on one side of the nsp8 α-helix and have the ability to form interactions with either mainly nsp7 (partly nsp8) in the SARS-CoV nsp7+8 (2:2) heterotetramer T1, mainly nsp8 (partly nsp7) in the SARS-CoV nsp7+8 heterotetramer T2 or only nsp7 in the FIPV nsp7+8 (2:1) heterotrimer (Figure S 16 A-C). Due to its sequence conservation, it is unlikely that alone this motif at BS II has a decisive impact for heterotrimer or -tetramer formation.

Therefore, unique interactions could exist, which explain the shift in complex stoichiometry from heterotrimer to-tetramer observed in the different CoVs categorized into group A, AB and B (Figure 4 B). Here, we identify a possibly heterotetramer stabilizing contact site in T2, where nsp8 Glu77 self-interacts with nsp8II Glu77, which gives the complex density and compactness (Figure 4 C). This residue is only present in nsp8 of SARS-CoV and SARS-CoV-2 from group A and MERS-CoV of group AB. However, homology models suggest that in the other tetramer forming complexes of group AB, HCoV-229E and PEDV, nsp8 Asn78 could partially replace this interaction (Figure 4 D). This is different in group B viruses, forming only heterotrimers, where residues at these positions are nsp8 Val77 and Asp78, with the Asp78 possibly being solvent exposed and hence unable to replace this interaction. Furthermore, we also identify a contact site possibly stabilizing the heterotrimer in the crystal structure of the FIPV nsp7+8 (2:1), which reveals that a second subunit of nsp7 (nsp7II) is locked via Phe76 to nsp8 (Figure 4 E). Importantly, this residue is uniquely conserved among trimer forming complexes of group B and group AB, but replaced by nsp7 Leu76 in the strictly heterotetramer forming group A.

These findings are compared to the recently released structure of the polymerase complex (pdb 6xez, Figure S 16 D and E), comprising nsp7+8+12+13(1:2:1:2) [29]. The residues potentially responsible for a shift in quaternary structure, nsp8 Glu77 or Asp78 and nsp7 Phe76, are distant from any protein-protein or protein-RNA interaction and thus are not expected to play a role in polymerase complex formation. Surprisingly, the identical set of residues in BS II supports all interactions (Glu60, Met62, Ala63, Met67 and Met70) between nsp8b and nsp12/nsp13.1 and between nsp8a and nsp13b (Figure S 16 D and E). Notably, within the polymerase complex amino acids involved in RNA binding point in the opposite direction of the protein interfaces and have little or no role in nsp7+8 complex formation.

## Discussion

Our findings reveal the nsp7+8 quaternary composition of seven CoVs representing five coronavirus species of the genera *Alpha*- and *Betacoronavirus*. Viruses of the same species (SARS-CoV/SARS-CoV-2 and TGEV/FIPV, respectively) produce the same type of nsp7+8 complexes. Next to a conserved nsp7+8 heterodimer (1:1), the inherent specificity of nsp7+8 complex formation categorizes them into three groups: group A forming only heterotetramers (2:2), group B forming only heterotrimers (2:1) and group AB forming both heterotetramers (2:2) and heterotrimers (2:1). Complexes of the same stoichiometry exhibit a conserved topology, consisting of an nsp8 homodimeric scaffold for the heterotetramers and an nsp7 homodimeric core for the trimers. Candidate structures based on our results highlight *Alpha*- and *Betacoronavirus*-wide conserved binding sites on nsp8, named BS I and BS II, which provide the modular framework for a variety of complexes. Furthermore, unique molecular contacts for the complex groups have the potential to determine the ability and preference for heterotrimer and/or heterotetramer formation.

We provide evidence that, even at high concentrations, the SARS-CoV-2 nsp7+8 heterotetramer (2:2) represents the predominant species. In order to relate our results to *in vivo* conditions, we consider the following aspects: According to maximum molecular crowding [30], polyproteins pp1a and pp1ab can reach a maximum of 125-450 μM, which translates to 3.9-11.7 mg/mL nsp7+8. This range is covered by our DLS and SAXS analysis. In absence of other interaction partners, we expect that, *in vivo*, the nsp7+8 (2:2) heterotetramer represents the predominant nsp7+8 complex of SARS-CoV-2 and other heterotetramer forming CoVs of complexation groups A and AB.

The heterotetramer candidate structures and models presented here are based on the conformers T1 or T2 of the SARS-CoV heterohexadecamer structure, which contains full-length nsp8 [20]. Although, our results cannot clarify, if one of these conformers is the biologically relevant structure existing in solution, the combined evidence provided here strongly suggests structural similarity to T1/T2. Considering the crystallographic origin of T1/T2 and the overlap of binding sites, the heterotetramer could well be a flexible and dynamic structure in solution.

In contrast to our findings, a SARS-CoV nsp7+8 hexadecamer structure has been reported [20]. However, this structure has been derived from X-ray crystallography, hence showing a static, frozen state, where the crystal lattice formation favors stabilized arrangements that could differ from the solution state of the protein complexes. In the case of nsp8, the flexible *N*-terminus could inhibit crystal formation, and has been removed in some studies [22, 23]. Alternatively, it may stabilize specific interactions, thereby promoting crystal formation by binding to one of the multiple interfaces presented between nsp7 and nsp8, resulting in a physiologically irrelevant larger oligomeric structure. The SAXS data presented here partially supports this scenario at high protein concentrations but also confirms a predominantly heterotetrameric assembly in solution. Thus, a potential shift of quaternary structure from heterotetramer towards a higher-order complex, such as a heterohexadecamer appears unlikely unless triggered e.g. by binding to nucleic acids as has been repeatedly described for nsp7+8 complexes [20].

All seven CoV nsp7 and nsp8 proteins shown here also form heterodimers (1:1). Such heterodimeric subcomplexes with nsp7 bound to nsp8 BS I have been observed in all deposited complex structures containing nsp7+8 [20–23] or nsp12 [13–16]. Therefore, the heterodimer represents the most basic form of nsp7+8 complexes and likely serves as universal substructure building block in the coordinated assembly of functional RTCs of CoVs from the genera *Alpha-* and *Betacoronavirus*. Moreover, heterotrimer and -tetramer formation are based on a second canonical binding site at the nsp8 *N*-terminal domain, BS II. This site appears to have a high propensity to form complexes with various binding partners (e.g. nsp7+8, nsp12 or nsp13). Accordingly, our analysis suggests that the nsp8 BS II strives for occupation. The nsp7+8 quaternary composition, topology and analysis of binding sites presented here allow us to reconstruct and propose a model of the complex formation pathway (Figure 5).

**Figure 5:**
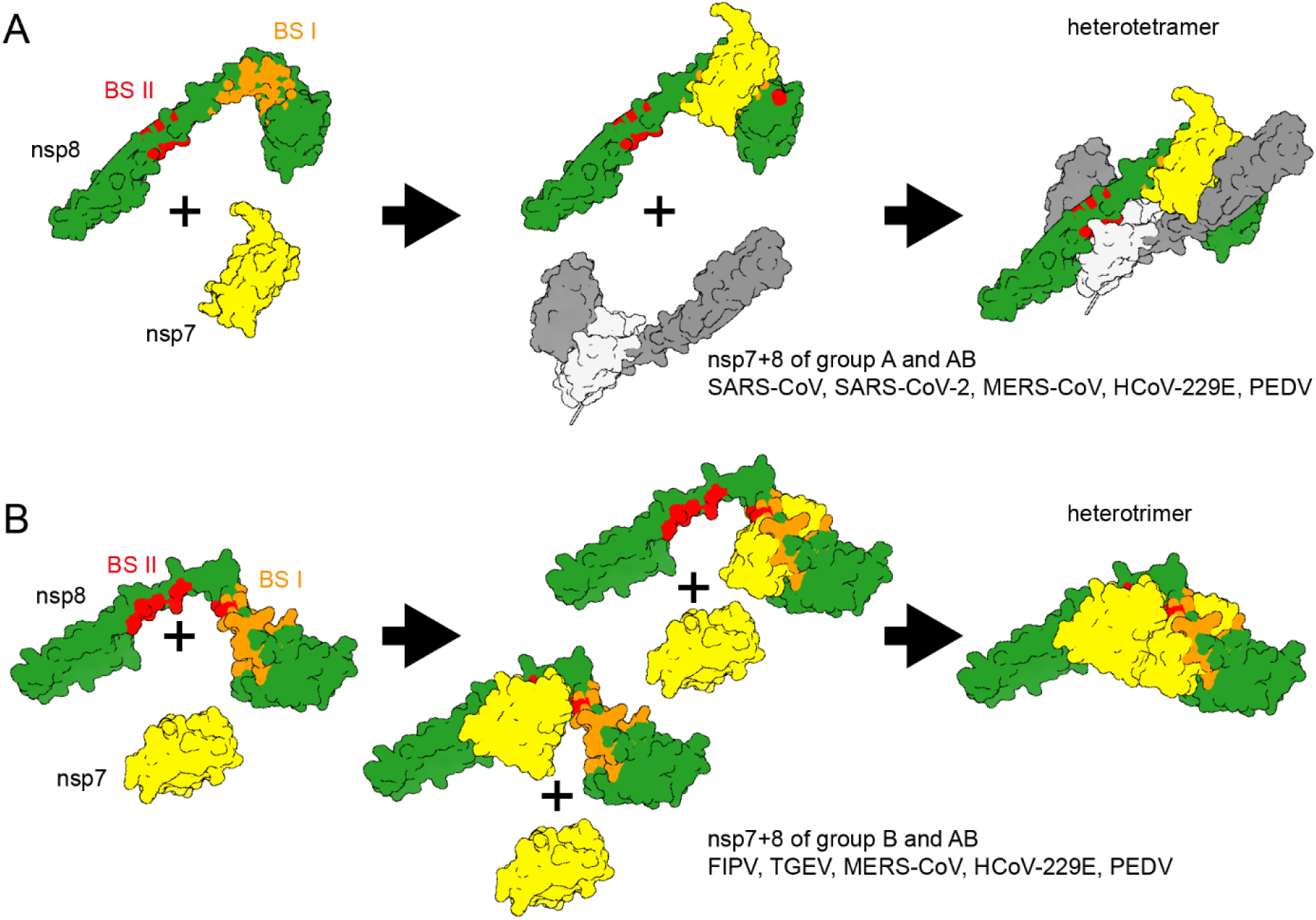
Proposed model for nsp7+8 complex formation. (**A**) For complexes of group A, heterodimers form via nsp8 BS I, which quickly dimerize via BS II into a heterotetramer. A theoretic route via a preformed nsp8 scaffold is unlikely to play a role in heterotetramer formation since no nsp7+8 (1:2) intermediates are observed for complexes of SARS-CoV or SARS-CoV-2. Moreover, nsp7 and nsp8 occupy neighboring positions in the replicase polyproteins, thus favoring their interaction (*in cis*) at early stages in the infection cycle (when intracellular viral polyprotein concentrations are low) over intermolecular interactions between different replicase polyprotein molecules as is also evident from the low dimerization ability of the precursors. (**B**) For group B complexes, we propose the formation of a heterodimer intermediate via nsp8 BS I or BS II and subsequent recruitment of a second nsp7, resulting an nsp7+8 (2:1) heterotrimer. This is also supported by the relatively high peak fractions of heterodimers detected. Group AB complexes can use both complexation pathways. In line with this, the proteins also produce a relatively high heterodimer signal but, ultimately, prefer to form heterotetramers rather than heterotrimers.

The preference for heterotrimer and -tetramer can probably be pinpointed to just a few amino acids within nsp8 BS II or interacting with it. Here, we identify two contacts that could have unique discriminatory potential for promoting heterotrimeric (nsp7 Phe76) or heterotetrameric (nsp8 Glu77 and Asn78) quaternary structures. Notably in presence of nsp7 Phe76 and nsp8 Asn78, as observed for group AB, the heterotetramer is always more abundant than the heterotrimer. However, compared to the entire BS II, theses contacts only represent a small share of the binding interface and contribute little interaction energy through van der Waals forces. Nevertheless, the unique position of their contacts could critically determine the types of interactions with one or another binding partner.

Since the critical residues required for nsp7+8 complex formation have no overlap with nsp12 interaction sites, direct docking of preformed heterotrimers and -tetramers to nsp12 can be expected. Furthermore, heterotrimeric and -tetrameric structures are compatible with accommodation of specific RNA structures similar to what has been suggested for heterohexadecameric nsp7+8 by Rao et al. [14]. Notably, if heterotrimeric or -tetrameric nsp7+8 structures were associated with nsp12, the binding site for nsp13 would be blocked, which may have regulatory implications for CoV replication. Altogether, these conserved binding mechanisms and overlapping binding sites confirm the proposed role of nsp8 as a major interaction hub within the CoV RTC [31], and indicate critical regulatory functions by specific nsp7+8 complexes.

Finally, we can only speculate about possible reasons for the existence of different nsp7+8 complexes: (1) Similar kinetic stability due to occupation of both binding sites (both structures exist because they are equally efficient in occupying BS I and BS II), (2) unknown functional relevance in CoV replication (e.g. specificity to RNA structures channeled to the nsp12 RdRp) or (3) adaption to host factors and possible regulatory functions.

In summary, our work shows, and provides a framework to understand, the characteristic distribution and structures of nsp7+8 (1:1) heterodimer, (2:1) heterotrimers and (2:2) heterotetramers in representative alpha- and betacoronaviruses. The nsp7+8 structure in solution can be used to investigate its independent functional role in the formation of active polymerase complexes and, possibly, regulation and coordination of polymerase and other RTC activities, for example in the context of antiviral drug development targeting different subunits of CoV polymerase complexes reconstituted *in vitro*.

## Material and Methods

### Cloning and gene constructs

The codon optimized sequence for the SARS-CoV-2 nsp7-8 region (NC_045512.2) was synthesized by Eurofins scientific SE with overhangs suitable for insertion into pASK-IBA33plus plasmid DNA (IBA Life Sciences). A golden gate assembly approach using Eco31I (BsaI) (Thermo Fisher Scientific) was used to shuttle the gene into the plasmid. Linker and tag of the expression construct SARS-CoV-2 nsp7-8-His_6_ contained the *C*-terminal amino acids–SGSGSARGS-His_6_ (SGSG residues as P1’-P4’ of Mpro cleavage site and SARGS-His_6_ residues as the default linker of pASK vectors). The SARS-CoV nsp7-8 pASK33+ plasmid generated previously [21, 32] was used for the expression of SARS-CoV nsp7-8-His_6_ containing the *C*-terminal amino acids–SARGS-His_6_. The expression plasmid for SARS-CoV M^pro^ was generated as described by Xue et al.[33]. To produce nsp7-8-SGSGSARGS-His_6_ precursor proteins in *Escherichia coli*, the nsp7-8 coding sequences of HCoV-229E (HCoV-229E; GenBank accession number AF304460), FIPV (FIPV, strain 79/1146; DQ010921), SARS-CoV (strain Frankfurt-1; AY291315), PEDV (PEDV, strain CV777, NC_003436) and TGEV (TGEV, strain Purdue; NC_038861) were amplified by reverse transcription-polymerase chain reaction (RT-PCR) from viral RNA isolated from cells infected with the respective viruses and inserted into pASK3-Ub-CHis_6_ using restriction- and ligation-free cloning methods as described before (Tvarogová et al., 2019). Similarly, the nsp7-9 or nsp7-11 coding region of MERS-CoV (strain HCoV-EMC; NC_019843), was amplified by RT-PCR from infected cells and inserted into pASK3-Ub-CHis_6_. The HCoV-229E and FIPV nsp5 coding sequences were cloned into pMAL-c2 plasmid DNA (New England Biolabs) for expression as MBP fusion proteins containing a *C*-terminal His_6_-tag. Primers used for cloning and mutagenesis are available upon request.

### Expression and purification

SARS-CoV M^pro^ was produced with authentic ends as described in earlier work [33]. To produce the precursors, SARS-CoV and SARS-CoV-2 nsp7-8-His_6_, BL21 Rosetta2 (Merck Millipore) were transformed, grown in culture flasks to OD_600_ = 0.4-0.6, then induced with 50 μM anhydrotetracycline and continued to grow at 20 °C for 16 h. For pelleting, cultures were centrifuged (6000 × *g* for 20 min) and cells were frozen at −20 °C. Cell pellets were lysed in 1:5 (*v*/*v*) buffer B1 (40 mM phosphate buffer, 300 mM NaCl, pH 8.0) with one freeze-thaw cycle, sonicated (micro tip, 70 % power, 6 times on 10 s, off 60 s; Branson digital sonifier SFX 150) and then centrifuged (20,000 × *g* for 45 min). Proteins were isolated with Ni^2+^-NTA beads (Thermo Fisher Scientific) in gravity flow columns (BioRad). Proteins were bound to beads equilibrated with 20 column volumes (CV) B1 + 20 mM imidazole, then washed with 20 CV B1 + 20 mM imidazole followed by 10 CV of B1 + 50 mM imidazole. The proteins were eluted in eight fractions of 0.5 CV B1 + 300 mM imidazole. Immediately after elution, fractions were supplemented with 4 mM DTT. Before analysis with native MS, Ni^2+^-NTA eluted fractions containing the polyprotein were concentrated to 10 mg/mL and further purified over a 10/300 Superdex 200 column (GE healthcare) in 20 mM phosphate buffer, 150 mM NaCl, 4 mM DTT, pH 8.0. The main elution peaks contained nsp7-8. For quality analysis, SDS-PAGE was performed to assess the sample purity.

To obtain a pre-purified SARS-CoV-2 nsp7+8 complex for DLS and SAXS, eluate fractions from the Ni^2+^-NTA column containing the nsp7-8-His_6_ were concentrated and the buffer exchanged with a PD-10 desalting column (GE Healthcare) equilibrated with 50 mM Tris, pH 8.0, 100 mM NaCl, 4 mM DTT and 4 mM MgCl_2_ (SEC-buffer). Then nsp7-8-His_6_ was eluted with 3.5 mL SEC-buffer and subsequently cleaved with M^Pro^-His_6_ (1:5, M^Pro^-His_6_: nsp7-8-His_6_) for 16 h at RT. M^pro^-His_6_ was removed with Ni-NTA agarose and the cleaved nsp7+8 complex was subjected to a HiLoad 16/600 Superdex 75 pg size exclusion column equilibrated with SEC-buffer.

The HCoV-229E, PEDV, FIPV and TGEV nsp7-8-His_6_ and MERS-CoV nsp7-11-His_6_ precursor proteins were produced and purified as described before (Tvarogová et al., 2019) with a slightly modified storage buffer. Anion exchange chromatography fractions of the peak containing the desired protein were identified by SDS-PAGE, pooled and dialyzed against storage buffer (50 mM Tris-Cl, pH 8.0, 200 mM NaCl and 2 mM DTT).

MBP-nsp5-His_6_ fusion proteins were purified using Ni^2+^-IMAC as described before (Tvarogová et al., 2019). To produce HCoV-229E and FIPV MBP-nsp5-His_6_, *E. coli* TB1 cells were transformed with the appropriate pMAL-c2-MBP-nsp5-His_6_ construct and grown at 37 °C in LB medium containing ampicillin (100 μg/mL). When an OD_600_ of 0.6 was reached, protein production was induced with 0.3 mM isopropyl β-D-thiogalactopyranoside (IPTG) and cells were grown for another 16 h at 18 °C. Thereafter, the cultures were centrifuged (6000 × *g* for 20 min) and the cell pellet was suspended in lysis buffer (20 mM Tris-Cl, pH 8.0, 300 mM NaCl, 5 % glycerol, 0.05 % Tween-20, 10 mM imidazole and 10 mM β-mercaptoethanol) and further incubated with lysozyme at 4 °C (0.1 mg/mL) for 30 min. Subsequently, cells were lysed by sonication and cell debris was removed by centrifugation for 30 min at 40,000 × *g* and 4°C. The cell-free extract was bound to pre-equilibrated Ni^2+^-NTA (Qiagen) matrix for 3 h. Ni^2+^-IMAC elution fractions were dialyzed against buffer comprised of 20 mM Tris-Cl, pH 7.4, 200 mM NaCl, 5 mM CaCl_2_ and 2 mM DTT and cleaved with factor Xa to release nsp5-His_6_. Then, nsp5-His_6_ was passed through an amylose column and subsequently bound to Ni^2+^-NTA matrix to remove any remaining MBP. Following elution from the Ni^2+^-NTA column, nsp5-His_6_ was dialyzed against storage buffer (20 mM Tris-Cl, pH 7.4, 200 mM NaCl and 2 mM DTT) and stored at −80 °C until further use.

### Native mass spectrometry

To prepare samples for native MS measurements, M^pro^ was buffer exchanged into 300 mM AmAc, 1 mM DTT, pH 8.0 by two cycles of centrifugal gel filtration (Biospin mini columns, 6,000 MWCO, BioRad) and the precursors were transferred into 300 mM AmAc, 1 mM DTT, pH 8.0 by five rounds of dilution and concentration in centrifugal filter units (Amicon, 10,000 MWCO, Merck Millipore). Cleavage and complex formation was started by mixing nsp7-8-His_6_ and protease M^pro^ with final concentrations of 15 μM and 3 μM, respectively. Three independent reactions were started in parallel and incubated at 4 °C overnight.

Tips for nano-electrospray ionization (nanoESI) were pulled in-house from borosilicate capillaries (1.2 mm outer diameter, 0.68 mm inner diameter, with filament, World Precision Instruments) with a micropipette puller (P-1000, Sutter Instruments) using a squared box filament (2.5 × 2.5 mm, Sutter Instruments) in a two-step program. Subsequently, tips were gold-coated using a sputter coater (Q150R, Quorum Technologies) with 40 mA, 200 s, tooling factor 2.3 and end bleed vacuum of 8×10^-2^ mbar argon.

Native MS was performed at a nanoESI quadrupole time-of-flight (Q-TOF) instrument (Q-TOF2, Micromass/Waters, MS Vision) modified for higher masses [34]. Samples were ionized in positive ion mode with voltages applied at the capillary of 1300-1500 V and at the cone of 130-135 V. The pressure in the source region was kept at 10 mbar throughout all native MS experiments. For desolvation and dissociation, the pressure in the collision cell was 1.5 × 10^-2^ mbar argon. For native MS, accelerating voltages were 10 - 30 V and quadrupole profile 1,000 - 10,000 *m*/*z*. For CID-MS/MS, acceleration voltages were 30 - 200 V. Raw data were calibrated with CsI (25 mg/mL) and analyzed using MassLynx 4.1 (Waters). Peak deconvolution and determination of relative intensity was performed using UniDec [35]. All determined masses are provided (Table S 1).

### XL-MALDI

Pre-purified FIPV nsp7+8 and HCoV-229E nsp7+8 at 20 μM were cross-linked with 0.15 % glutaraldehyde (Sigma-Aldrich) at 4 °C for 25 min before diluting them to 1 μM in MALDI matrix solution (sinapinic acid 10 mg/mL in acetonitrile/water/TFA, 49.95/49.95/0.1, *v*/*v*/*v*) and spotting (1 μL) onto a stainless steel MALDI target plate. The MALDI-TOF/TOF mass spectrometer (ABI 4800, AB Sciex) equipped with a high-mass detector (HM2, CovalX) was used in linear mode. For acquiring mass spectra (1,000 to 1,000,000 *m/z*) spots were ionized with a Nd:YAG laser (355 nm) and 500 shots per spectrum were accumulated. Obtained raw data were smoothed and analyzed using mMass (v5.5.0, by Martin Strohalm [36]).

### DLS

To check the monodispersity of the samples and to study the stoichiometry of the nsp7+8 complexes, DLS measurements were performed with the Spectro Light 600 (Xtal Concepts). The complex was concentrated to various concentrations and samples were spun down for 10 min at 12,000 rpm and 4 °C. A Douglas Vapour batch plate (Douglas instruments) was filled with paraffin oil and 2 μL of each sample was pipetted under oil. DLS measurements for each sample were performed at 20 °C with 20 measurements for 20 s each, respectively.

### SAXS

SAXS data were collected on the P12 beamline of EMBL at the PETRA III storage ring (DESY, Hamburg). X-ray wavelength of 1.24 Å (10 keV) was used for the measurements, scattered photons were collected on a Pilatus 6M detector (Dectris), with a sample to detector distance of 3 m. Data were collected on 22 concentrations ranging from 1.2 to 48 mg/mL nsp7+8 in 50 mM Tris, pH 8.0, 100 mM NaCl, 4 mM DTT and 4 mM MgCl_2_, pure buffer was measured between samples. For each data collection, 20 frames of 100 ms were collected. 2D scattering images were radially averaged and normalized to the beam intensity. The frames were compared using the program Cormap [37, 38], and only similar frames were averaged and used for further analysis to avoid possible beam-induced effects. Scattering collected on the pure buffer was subtracted from that of the sample and the resulting curves were normalized to the protein concentration to obtain the scattering of nsp7+8 complexes.

The data processing pipeline SASflow was used for data reduction and calculation of the overall SAXS parameters [38]. Molecular weights were inferred from different molecular calculation methods using a Bayesian assessment [39]. The program CRYSOL was used to compute the theoretical curves from the atomic structures [40]. Volume fractions of the components of the oligomeric mixtures were computed and fitted to the data using the program OLIGOMER [41]. The dimer of T1 was built by the program SASREFMX [42], which builds a dimeric model that fits best, in mixture with the monomeric T1, multiple scattering curves collected at different concentrations.

### Visualization

Molecular graphics and analyses were performed with UCSF ChimeraX, developed by the Resource for Biocomputing, Visualization, and Informatics at the University of California, San Francisco, with support from National Institutes of Health R01-GM129325 and the Office of Cyber Infrastructure and Computational Biology, National Institute of Allergy and Infectious Diseases.

## Acknowledgements

We would like to thank the XBI User consortium for providing the instrumentation and laboratories at the European XFEL.

## Author contributions

Conceptualization and methodology - B.K., C.S., C.U. G.B., R.M. and J.Z.; plasmid construction and protein production - B.K., C.S., G.B., L.B.; providing research materials C.U., K.L. and J.Z.; XL-MALDI-MS - B.K., M.K. and R.Z.; SAXS: C.B., D.S. Investigation - B.K., G.B., C.S., R.S. Discussion of results - B.K., C.S., C.U., K.L., R.M., G.B., J.Z.; Formal analysis and visualization - B.K.; Original Draft - B.K. and C.U.; Writing, review and editing - B.K. and C.U. with help from all authors.

## Supplement

**Table S 1:** Mass table. Data from at least three representative native mass spectra were analyzed with MassLynx 4.1 and UniDec. Mtheo is calculated based on amino acid sequence. Full width half-maximum (FWHM) values derived from main peaks, charge state given in parenthesis. Three data points are used for calculation of average (Avg) and standard deviation (SD) (only two for PEDV complexes).

| | Oligomer | $M_{\text{theo}}$ (Da) | Avg $M_{\text{exp}} \pm \text{SD}$ (Da) | Avg FWHM (Da) and charge state | nsp7+8 relative intensity fraction (%) |
| --- | --- | --- | --- | --- | --- |
| SARS-CoV-2 | nsp7+8 (1:1) | 31121 | 31140 ± 10 | 62 (11+) | 4 ± 2 |
|  | nsp7+8 (2:1) | - | - | - | - |
| | nsp7+8 (2:2) | 62242 | 62300 ± 20 | 155 (15+) | 96 ± 2 |
| SARS-CoV | nsp7+8 (1:1) | 31129 | 31123 ± 2 | 20 (11+) | 2.3 ± 0.2 |
|  | nsp7+8 (2:1) | - | - | - | - |
| | nsp7+8 (2:2) | 62258 | 62267 ± 2 | 65 (15+) | 97.6 ± 0.2 |
| MERS-CoV | nsp7+8 (1:1) | 30931 | 30954.0 ± 0.0 | 20 (10+) | 73 ± 11 |
| | nsp7+8 (2:1) | 40012 | 40025 ± 2 | 22 (11+) | 8 ± 1 |
| | nsp7+8 (2:2) | 61898 | 61920 ± 10 | 75 (14+) | 19 ± 11 |
| PEDV | nsp7+8 (1:1) | 30895 | 30927 ± 1 | 20 (10+) | 52 ± 4 |
| | nsp7+8 (2:1) | 40102 | 40238 ± 3 | 27 (11+) | 6 ± 1 |
| | nsp7+8 (2:2) | 61723 | 61863 ± 1 | 45 (15+) | 42 ± 2 |
| HCoV-229E | nsp7+8 (1:1) | 30901 | 30920 ± 10 | 20 (10+) | 20 ± 7 |
| | nsp7+8 (2:1) | 40188 | 40145 ± 1 | 29 (11+) | 12 ± 1 |
| | nsp7+8 (2:2) | 61793 | 61839 ± 1 | 45 (15+) | 69 ± 6 |
| TGEV | nsp7+8 (1:1) | 30930 | 30962 ± 6 | 22 (11+) | 17 ± 3 |
| | nsp7+8 (2:1) | 40401 | 40470 ± 20 | 68 (12+) | 83 ± 3 |
|  | nsp7+8 (2:2) | - | - | - | - |
| FIPV | nsp7+8 (1:1) | 308334 | 30850 ± 5 | 26 (11+) | 17 ± 2 |
| | nsp7+8 (2:1) | 40234 | 40400 ± 100 | 208 (12+) | 83 ± 2 |
|  | nsp7+8 (2:2) | - | - | - | - |

**Table S 2:** Sequence identity matrix of nsp7-8 species. Values for pairwise sequence identity are given in percent. For the multiple sequence alignment with identity matrix output the SIAS Sequence identity and similarity tool has been used provided by Secretaria general de sciencia, technologica e innovacion of Spain (http://imed.med.ucm.es/Tools/sias.html). As input parameter, length of the smallest sequence was selected.

|  |  |  |  |  |  |  |  |
| --- | --- | --- | --- | --- | --- | --- | --- |
| SARS-CoV-2 | 100 |  |  |  |  |  |  |
| SARS-CoV | 98 | 100 |  |  |  |  |  |
| HCoV-229E | 43 | 44 | 100 |  |  |  |  |
| PEDV | 42 | 42 | 71 | 100 |  |  |  |
| FIPV | 42 | 43 | 63 | 62 | 100 |  |  |
| TGEV | 42 | 42 | 64 | 61 | 94 | 100 |  |
| MERS | 28 | 28 | 21 | 21 | 20 | 20 | 100 |
| % | SARS-CoV-2 | SARS-CoV | HCoV-229E | PEDV | FIPV | TGEV | MERS |

**Table S 3:**
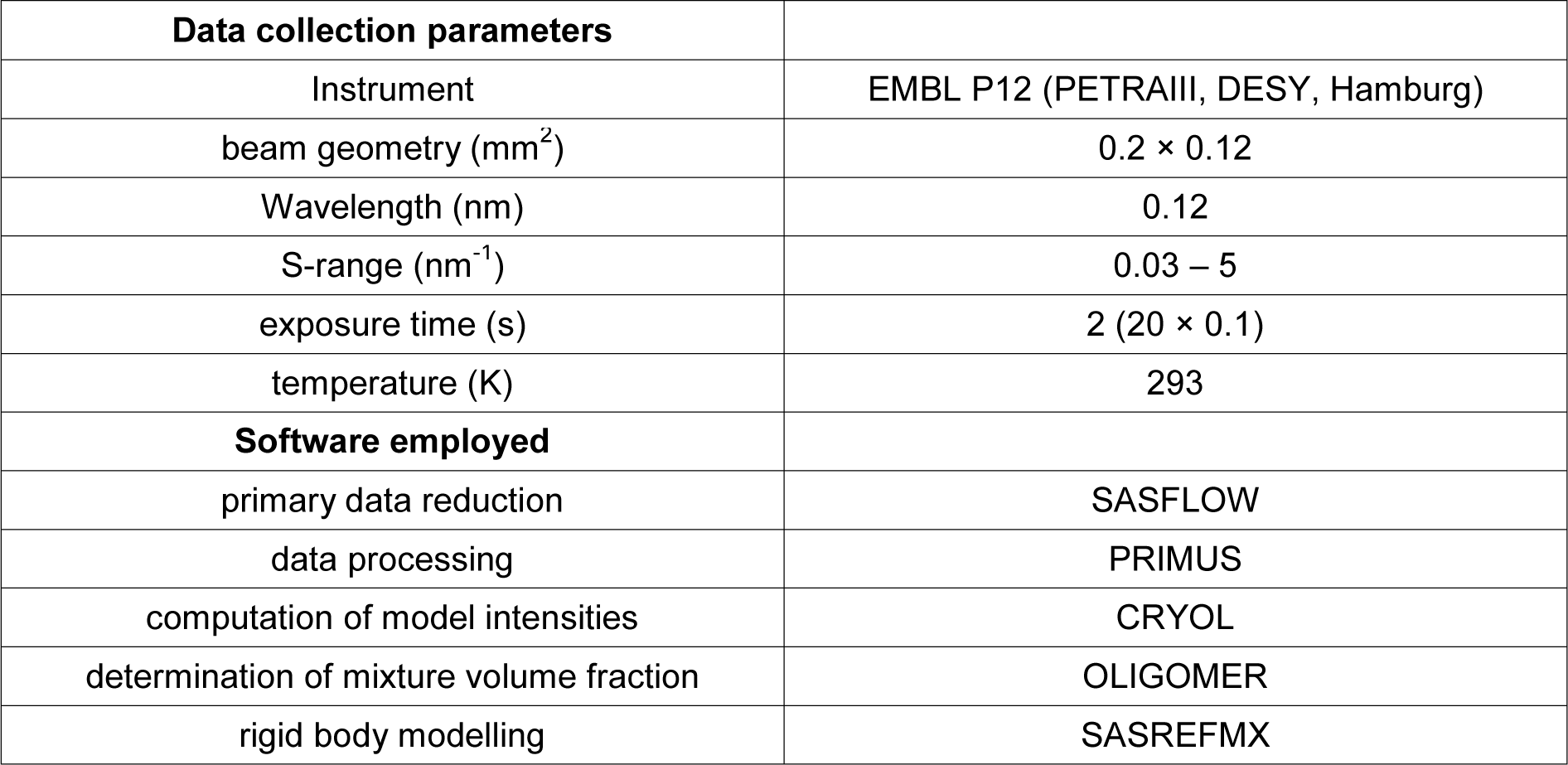
SAXS experimental parameters and analysis methods.

**Table S 4:** Overall parameters computed from the SAXS curves.

| c (mg/mL) | Rg (nm) | MW Bayes (kDa) | Fit to tetramer T1 | Mixture of tetramer and dimer of tetramer |  | Mixture of tetramer and hexadecamer |  |
| --- | --- | --- | --- | --- | --- | --- | --- |
|  |  |  | χ<sup>2</sup> | χ<sup>2</sup> | Tetramer volume fraction | χ<sup>2</sup> | Tetramer volume fraction |
| 1.2 | 3.25 | 46 | 1.35 | 1.34 | 1 | 1.34 | 1 |
| 1.2 | 3.41 | 58 | 1.27 | 1.27 | 1 | 1.27 | 1 |
| 2.1 | 3.37 | 56 | 1.31 | 1.41 | 0.97 | 1.47 | 1 |
| 3.9 | 3.46 | 60 | 2.57 | 2.54 | 0.9 | 3.7 | 0.98 |
| 4.4 | 3.41 | 62 | 1.97 | 2.2 | 0.9 | 3.34 | 0.97 |
| 7.7 | 3.56 | 62 | 5.13 | 2.3 | 0.8 | 4.84 | 0.95 |
| 9.1 | 3.67 | 67 | 14.32 | 2.45 | 0.69 | 9.54 | 0.91 |
| 10.4 | 3.79 | 62 | 22.95 | 4.4 | 0.73 | 15.02 | 0.93 |
| 10.6 | 3.79 | 74 | 21.79 | 2.86 | 0.66 | 12.75 | 0.91 |
| 11.5 | 3.66 | 62 | 4.94 | 2.14 | 0.79 | 4.77 | 0.94 |
| 12.8 | 3.73 | 74 | 62.98 | 5.13 | 0.63 | 30.59 | 0.9 |
| 13.1 | 3.53 | 62 | 19.63 | 5.29 | 0.77 | 15.77 | 0.94 |
| 16.6 | 3.82 | 74 | 17.89 | 2.27 | 0.64 | 9.96 | 0.9 |
| 20 | 3.67 | 68 | 40.19 | 5.59 | 0.68 | 20.25 | 0.91 |
| 21.1 | 3.82 | 74 | 67.56 | 4.62 | 0.61 | 29.52 | 0.89 |
| 25.8 | 3.8 | 74 | 73 | 4.69 | 0.57 | 27.08 | 0.88 |
| 26.6 | 3.86 | 74 | 58.68 | 6.49 | 0.62 | 24.02 | 0.89 |
| 31.1 | 3.91 | 83 | 148.76 | 7.82 | 0.52 | 45.67 | 0.86 |
| 32.1 | 3.82 | 76 | 111.04 | 10.92 | 0.61 | 41.36 | 0.89 |
| 47.7 | 3.95 | 86 | 86.73 | 6.57 | 0.47 | 21.69 | 0.84 |

**Table S 5:** Results table, overview of main experimental results for nsp7+8 complexes of seven CoV species. Results from left to right: Categorization of complexes, quantitation of complexes by native MS (values are given in percent), identified homodimers nsp7_2_ and nsp8_2_, and dimerization of precursor (clearly assigned species indicated with “yes” and if no assignment possible with minus(-)), nsp8 and nsp7 possible mutations that shift heterotrimer to heterotetramer formation.

|  | Group | Complex<br>nsp7+8<br>(1:1) | Complex<br>nsp7+8<br>(2:1) | Complex<br>nsp7+8<br>(2:2) | Dimer<br>nsp7<br>(2) | Dimer<br>nsp8 (2) | Dimer<br>nsp7-8 | nsp8<br>Glu77/<br>Asn78 | nsp7<br>Phe76 |
| --- | --- | --- | --- | --- | --- | --- | --- | --- | --- |
| SARS-CoV-2 | A | 4 | - | 96 | yes | yes | yes | Glu77 | - |
| SARS-CoV | A | 2 | - | 98 | yes | yes | yes | Glu77 | - |
| MERS-CoV | AB | 73 | 8 | 19 | yes | - | *different precursor | Glu77 | Phe76 |
| HCoV-229E | AB | 20 | 12 | 69 | yes | - | yes | Asn78 | Phe76 |
| PEDV | AB | 52 | 6 | 42 | yes | - | yes | Asn78 | Phe76 |
| FIPV | B | 17 | 83 | - | yes | - | - | - | Phe76 |
| TGEV | B | 17 | 83 | - | yes | - | - | - | Phe76 |

**Table S 6:**
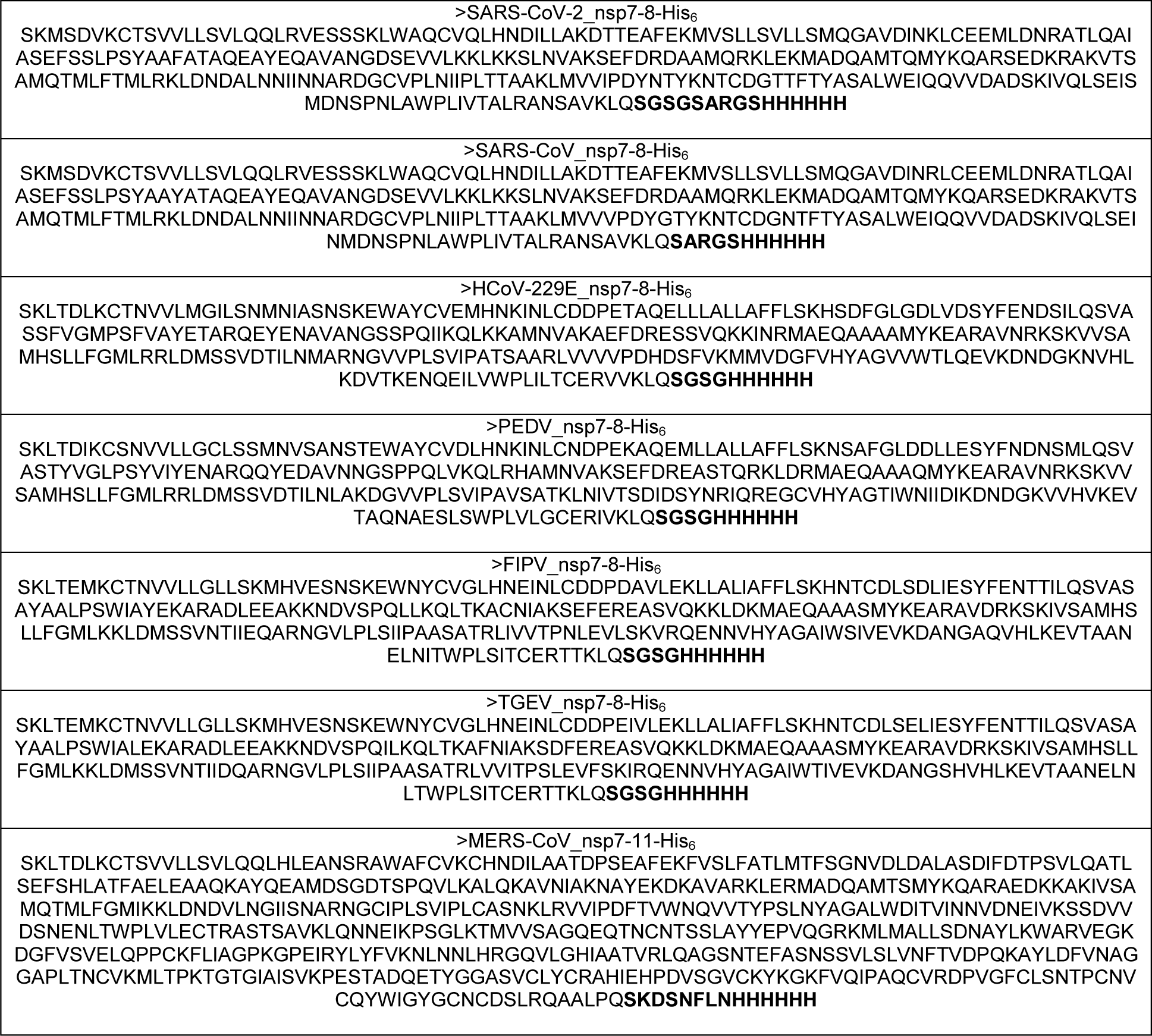
Amino acid sequences of precursor constructs.

**Figure S 1:**
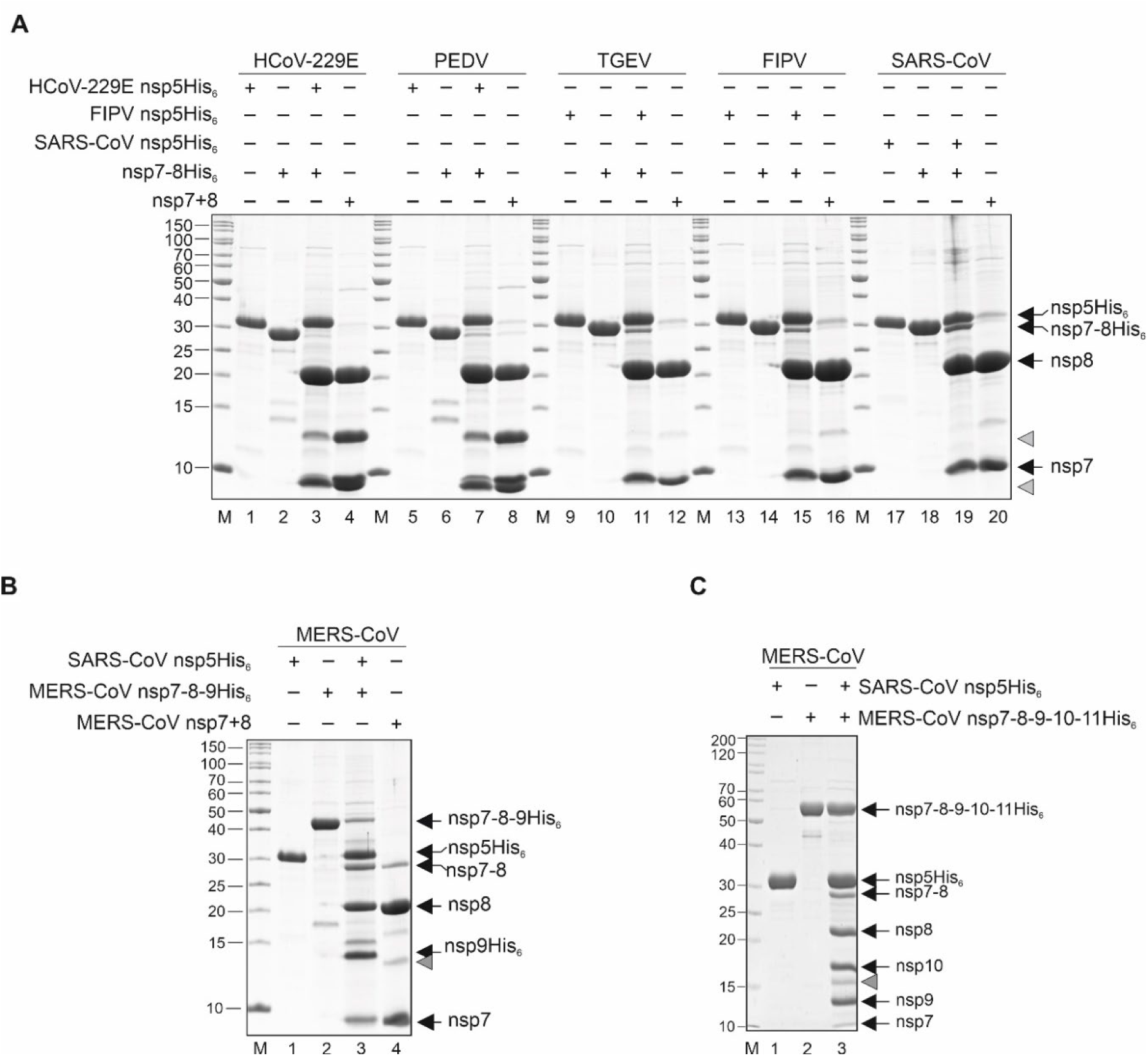
Mpro mediated processing of precursor protein constructs. SDS-PAGE analysis of M^pro^ (nsp5)-mediated processing and generation of coronavirus nsp7+8 complexes with authentic *N*- and *C*-termini from polyprotein precursors nsp7-8, nsp7-8-9 and nsp7-11. The nsp7+8 complexes of HCoV-229E, PEDV, FIPV, TGEV, and SARS-CoV were produced from the respective nsp7-8-His_6_ precursors and MERS-CoV was produced from an nsp7-9-His_6_ precursor. Precursor proteins were purified by Ni^2+^-IMAC and ion-exchange chromatography. Then, 15 μg protein was cleaved with M^pro^ (nsp5-His_6_, 5 μg) for 48 h at 4 °C. Subsequently, His_6_-tag containing cleavage products were removed by passing the material through a Ni^2+^-IMAC column and nsp7+8 complexes were enriched by ion-exchange chromatography. (**A**) SDS-PAGE showing the purified M^pro^ (nsp5-His_6_) – lanes 1, 5, 9, 13, 17; nsp7-8-His_6_ – lanes 2, 6, 10, 14, 18; M^pro^-mediated cleavage reaction – lanes 3, 7, 11, 15, 19; enriched nsp7+8 complexes – lanes 4, 8, 12, 16, 20. (**B**) SDS-PAGE showing the purified M^pro^ (nsp5-His_6_) – lane 1; nsp7-8-9-His_6_ – lane 2; M^pro^-mediated cleavage reaction – lane 3; enriched nsp7+8 complex – lane 4. (**C**) SDS-PAGE showing the purified M^pro^ (nsp5-His_6_) – lane 1; nsp7-8-9-10-11-His_6_ - lane 2; M^pro^ mediated cleavage reaction – lane 3. Lane M, marker proteins with molecular masses in kD indicated to the left. Black arrows on the right indicate the identities of proteins generated from precursor proteins by M^pro^-mediated cleavage. Gray arrowheads indicate aberrant *in vitro* cleavage products of nsp8 as observed previously for SARS-CoV [22]. nsp – nonstructural protein. +/- indicate the presence or absence of the respective proteins.

**Figure S 2:**
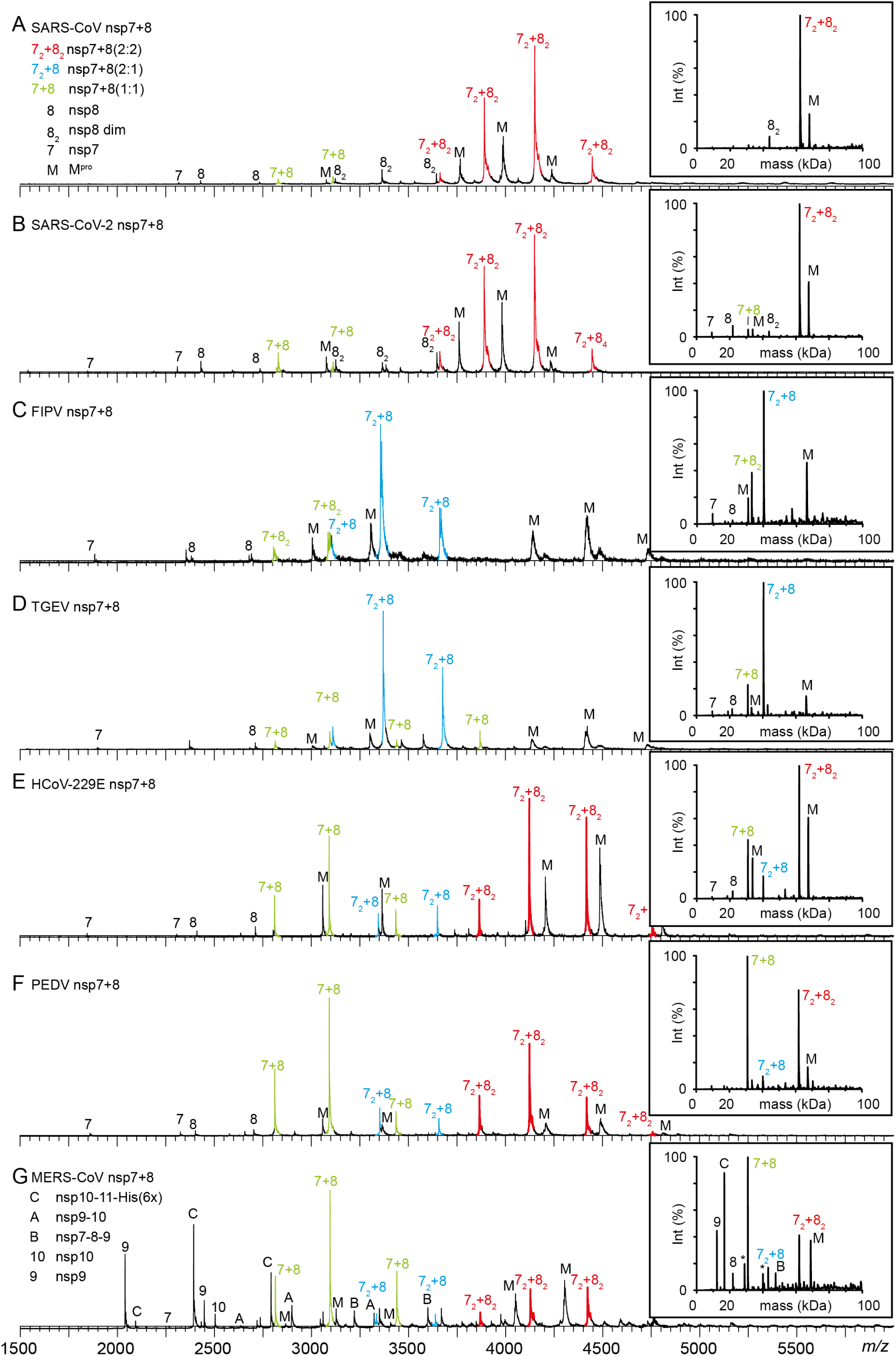
Native MS of nsp7+8 complexes of seven CoVs representing five different CoV species. Representative mass spectra showing distinct nsp7+8 complexation patterns that were classified into the three groups A, B and AB. Complex formation triggered by M^pro^ (M) mediated cleavage of 15 μM nsp7-8-His_6_ or MERS-CoV nsp7-11-His_6_ precursors in 300 mM AmAc, 1 mM DTT, pH 8.0. (**A**) SARS-CoV and (**B**) SARS-CoV-2 from group A forming nsp7+8 (2:2) heterotetramers (red), (**C**) FIPV and (**D**) TGEV from group B forming nsp7+8 (2:1) heterotrimers (blue) as well as (**E**) HCoV-229E and (**F**) PEDV from group AB forming both complex stoichiometries. (**G**) MERS-CoV, also from group AB, produced from an nsp7-11 precursor, additionally results in several processing intermediates that allow for an estimation of relative cleavage efficiencies at different cleavage sites. All groups form nsp7+8 (1:1) heterodimers as intermediate state (green).

**Figure S 3:**
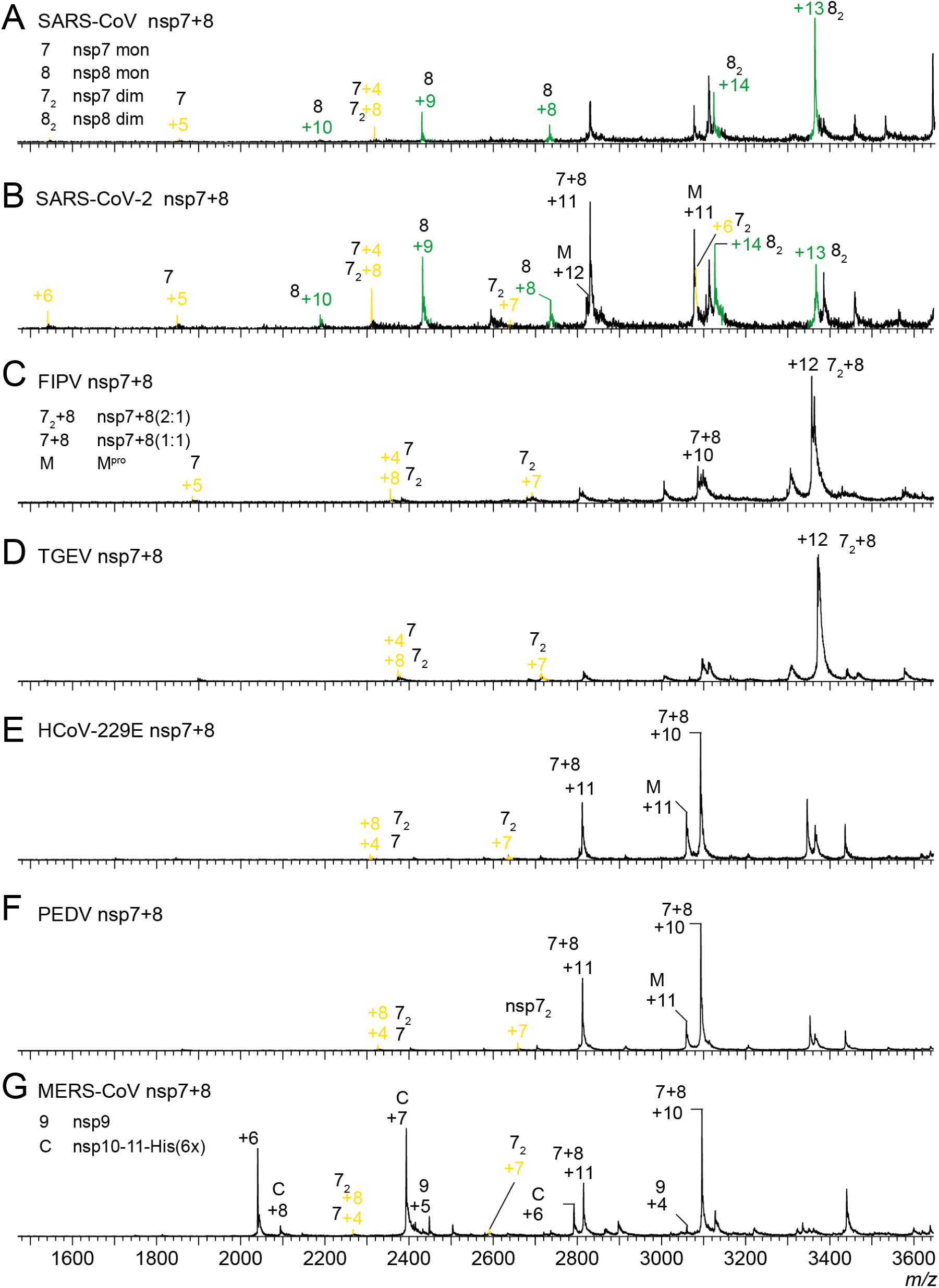
Zoom in on nsp7 and nsp8 monomers and homodimers from spectra in Figure S 2. Representative native mass spectra showing region 1500-3600 *m/z* (for full spectra see Figure S 2) of cleaved nsp7-8 or nsp7-11 precursors showing that homodimers of nsp8 (green, nsp8_2_) were detected for SARS-CoV (**A**) and SARS-CoV-2 (**B**), but not for (**C**) FIPV, (**D**) TGEV, (**E**) HCoV-229E, (**F**) PEDV and (**G**) MERS-CoV. Homodimers of nsp7 (yellow, nsp7_2_) were detected for all seven CoVs.

**Figure S 4:**
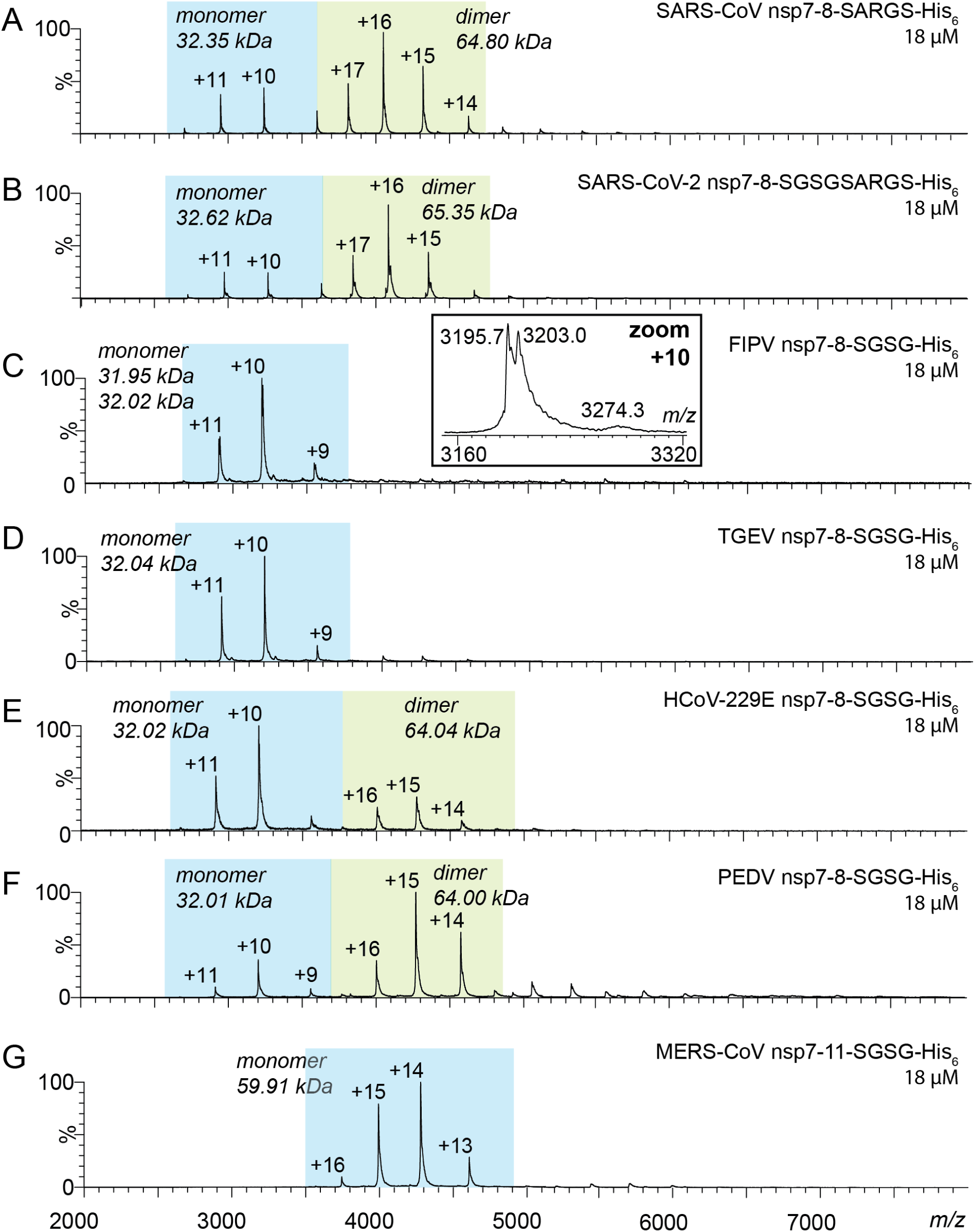
Mass and oligomeric state of nsp7-8 precursors. Native MS of nsp7-8 precursors (**A-G**) sprayed at 18 μM from 300 mM AmAc, pH 8.0 and 1 mM DTT. Dominant charge envelope is highlighted (blue box). Labelled are charge states and molecular mass. (**C**) Inset shows mass heterogeneity in FIPV nsp7-8. The experimental molecular weight *M_exp_* of the precursors agrees with the sequence-derived theoretical *M_theo_* (Table S 1). Only FIPV nsp7-8 contained two mass species separated by ~110 Da. This heterogeneity was attributed to the precursor’s central nsp8 domain following M^pro^ processing. Assignment to an amino acid variation failed but potentially was the result of codon heterogeneity in the plasmid. Nevertheless, both forms behaved identical and we refrained from further optimization.

**Figure S 5:**
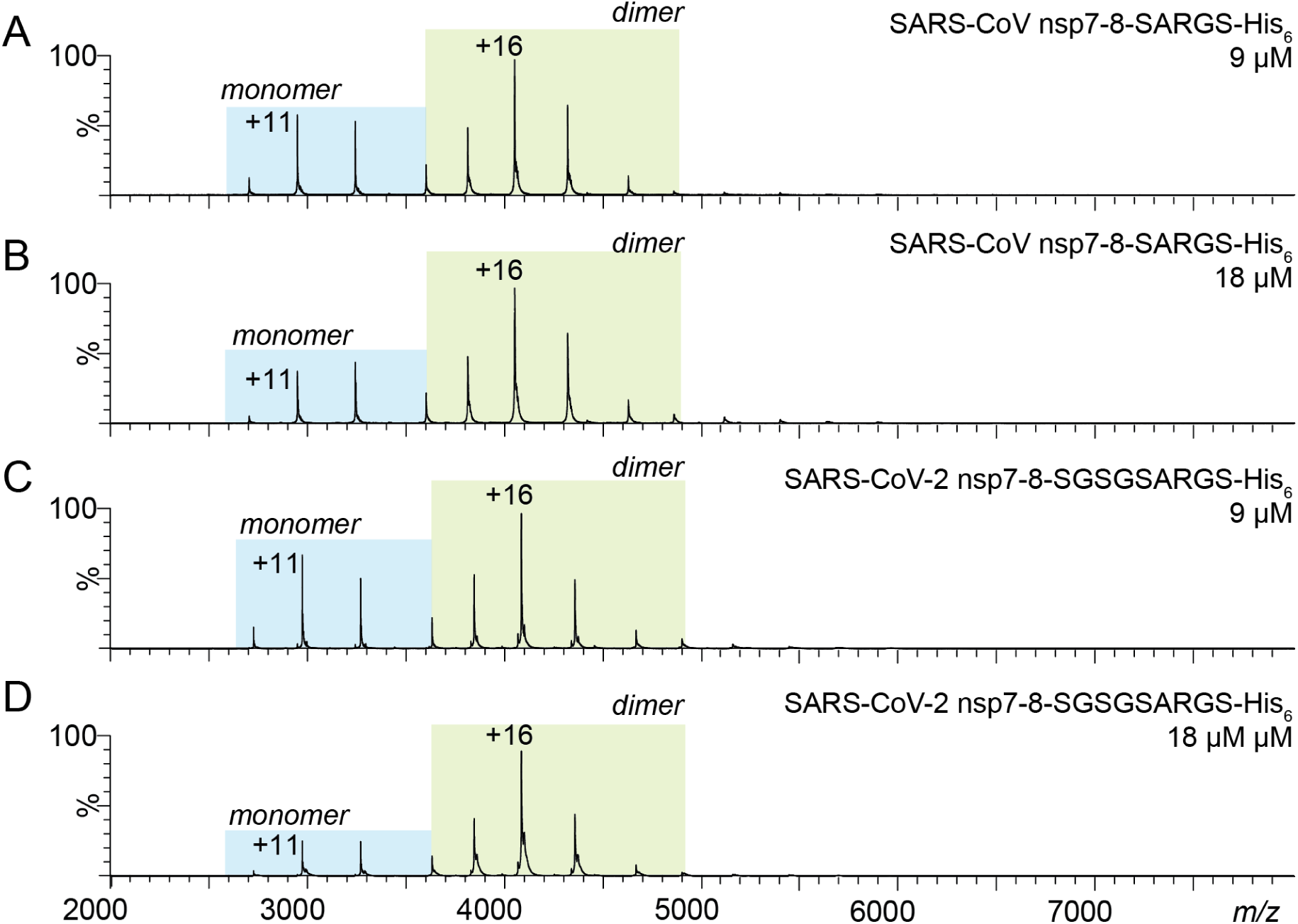
Monomer-dimer equilibrium in SARS-CoV and SARS-CoV nsp7-8 precursors. Native mass spectra of nsp7-8 precursors of SARS-CoV (**A**) 18 μM and (**B**) 9 μM and SARS-CoV-2 (**C**) 18 μM and (**D**) 9 μM. Proteins at 18 μM were diluted to 9 μM, incubated for 10 min and then sprayed from 300 mM AmAc, pH 8.0 and 1 mM DTT. Charge states are labelled.

**Figure S 6:**
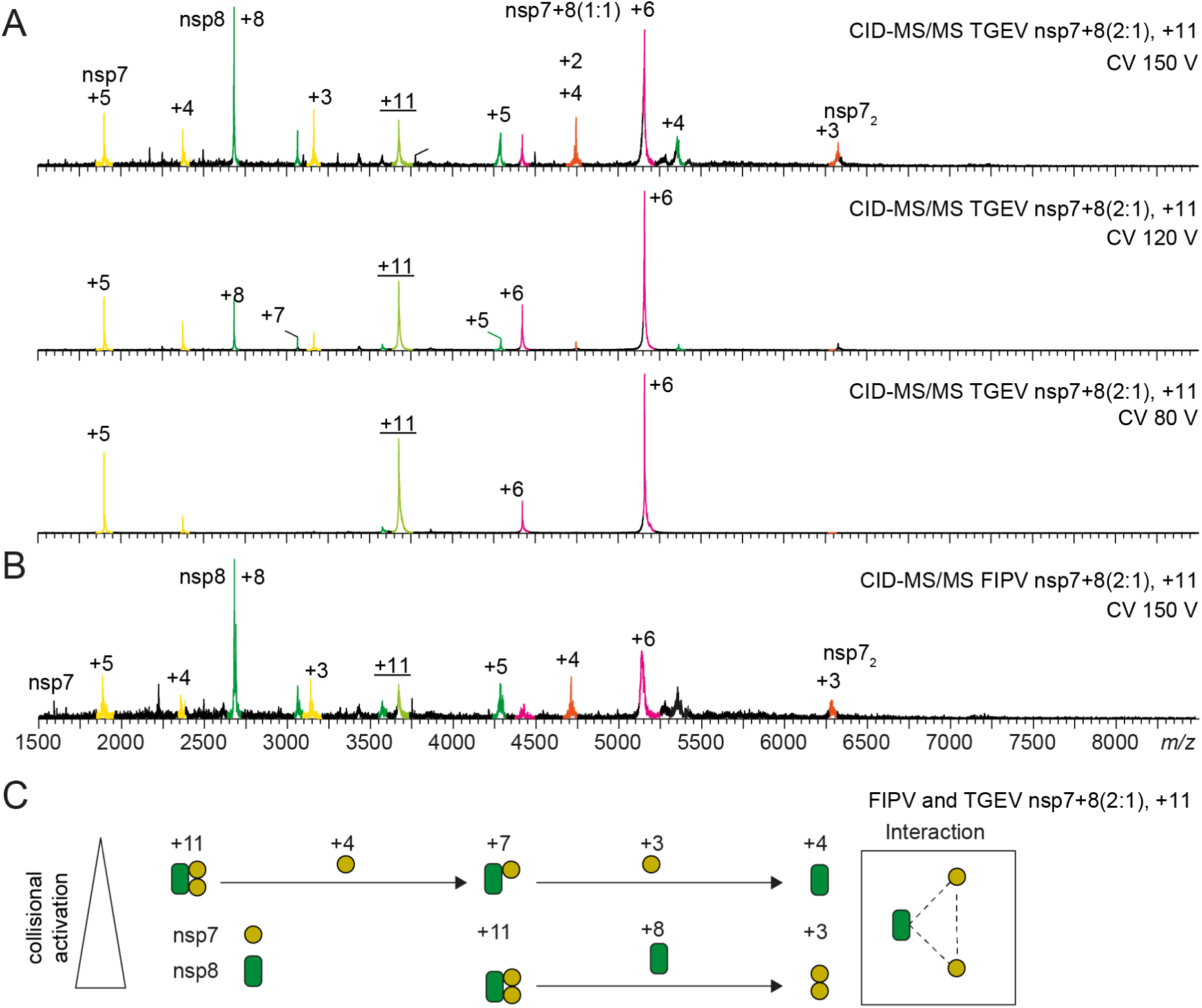
Topological reconstruction from CID-MS/MS of TGEV and FIPV nsp7+8 heterotrimers. (**A**) Three product ion spectra of TGEV 11+ nsp7+8 (2:1) heterotrimers are shown at different collisional voltages (CV). Increased collisional activation allows assignment of additional charge states and product species. (**B**) Product ion spectra of the FIPV 11+ nsp7+8 (2:1) show a similar dissociation pattern as its homologue from TGEV. (**C**) Schematic pathway of dissociation allows for a topological reconstruction of the TGEV and FIPV heterotrimers.

**Figure S 7:**
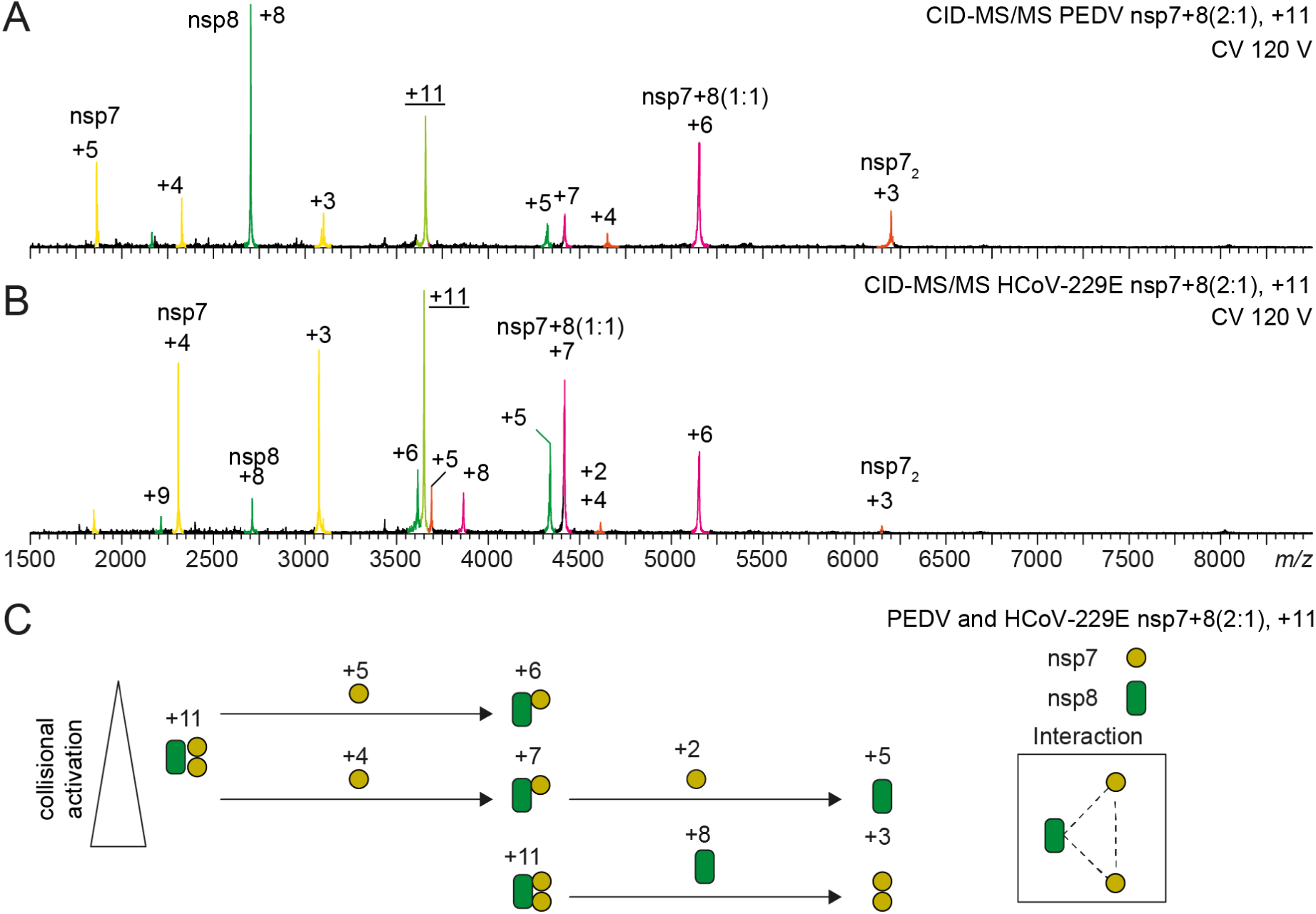
Topological reconstruction from CID-MS/MS of PEDV and HCoV-229E nsp7+8 heterotrimers. (**A**) Product ion spectra of the PEDV 11+ nsp7+8 (2:1) heterotrimer shows nsp7 homodimeric species, revealing the core interaction of this protein complex. (**B**) Product ion spectra of the HCoV-229E 11+ nsp7+8 (2:1) heterotrimer. Compared to the dissociation pattern of PEDV homologue, the HCoV-229E nsp7 and nsp8 high charge products have distinct intensities and charge state distributions but the detected product species are similar. (**C**) Schematic pathway of dissociation allows for a topological reconstruction of the PEDV and HCoV-229E heterotrimer.

**Figure S 8:**
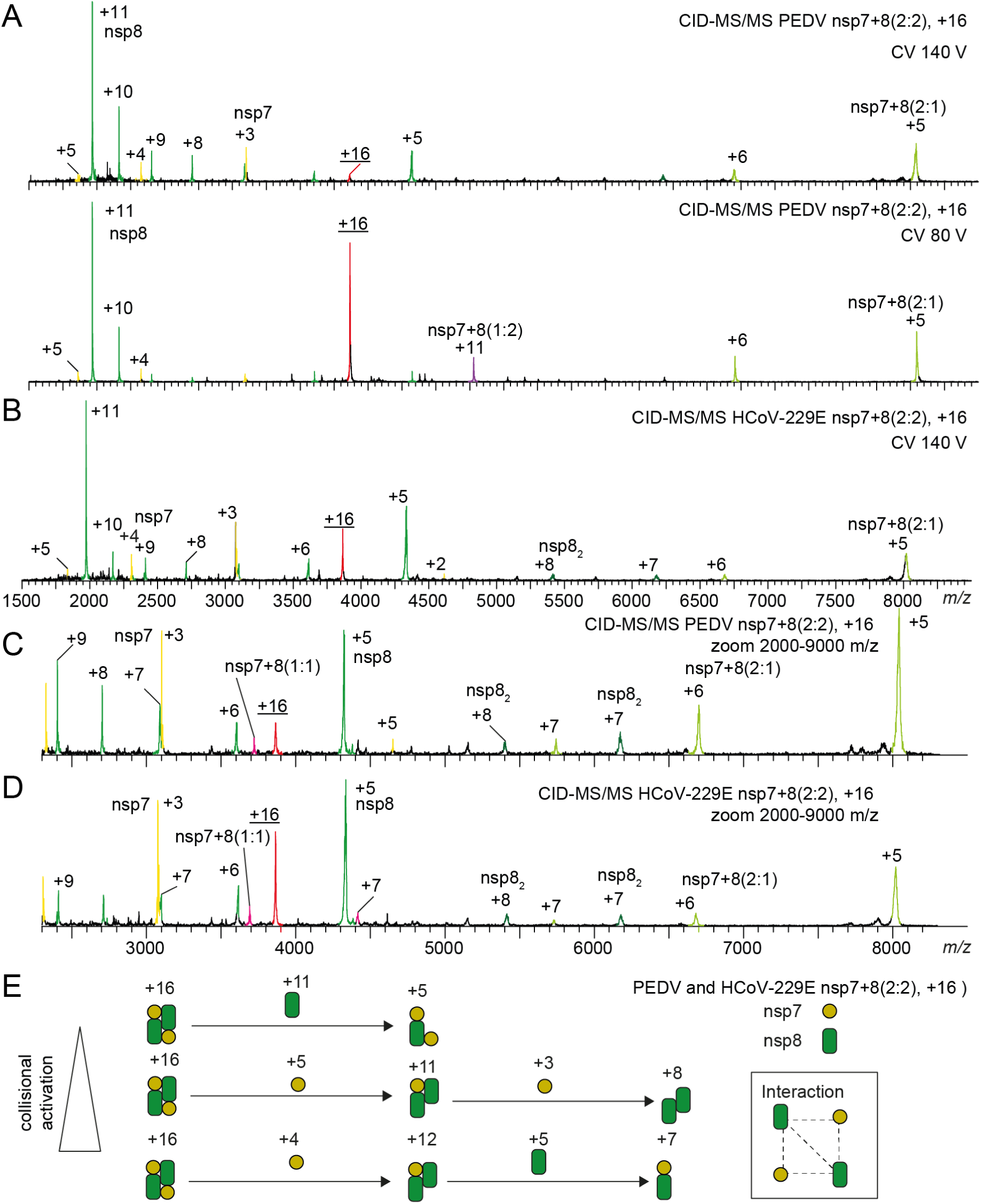
Topological reconstruction from CID-MS/MS of PEDV and HCoV-229E nsp7+8 heterotetramers. (**A**) Product ion spectra of PEDV 11+ nsp7+8 (2:1) heterotrimers shows nsp7 homodimeric species, revealing the core interaction of this protein complex. (**B**) Product ion spectra of HCoV-229E 11+ nsp7+8 (2:1) heterotrimers are shown. Compared to the dissociation pattern of PEDV homologue, the HCoV-229E nsp7 and nsp8 high charge products have distinct intensities and charge state distributions but the detected product species are similar. (**C**) Schematic pathway of dissociation allows for a topological reconstruction of the PEDV and HCoV-229E heterotrimer.

**Figure S 9:**
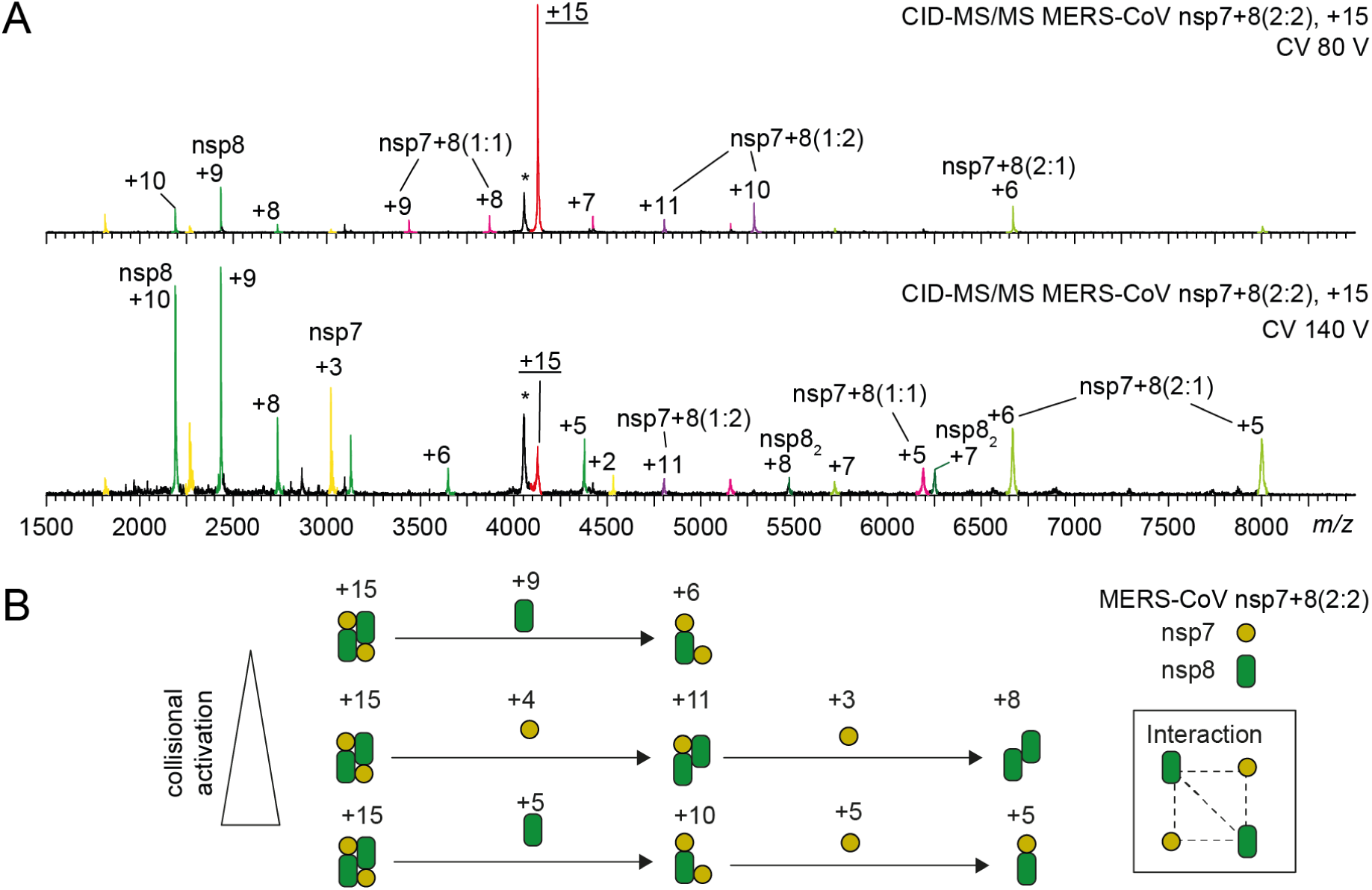
Topological reconstruction from CID-MS/MS of MERS-CoV nsp7+8 heterotetramers. (**A**) Product ion spectra of MERS-CoV 15+ nsp7+8 (2:2) heterotetramers are shown at two different CV. At 100 V CV, the dissociation of one nsp7 or, alternatively, one nsp8 occurs as indicated by the high and low charge products. At 140 V CV, other dissociation products increase in intensity (e.g. nsp8_2_ homodimer and nsp7+8(1:1) heterodimer). (**B**) Schematic pathway of dissociation allows topological reconstruction of the MERS-CoV heterotetramers.

**Figure S 10:**
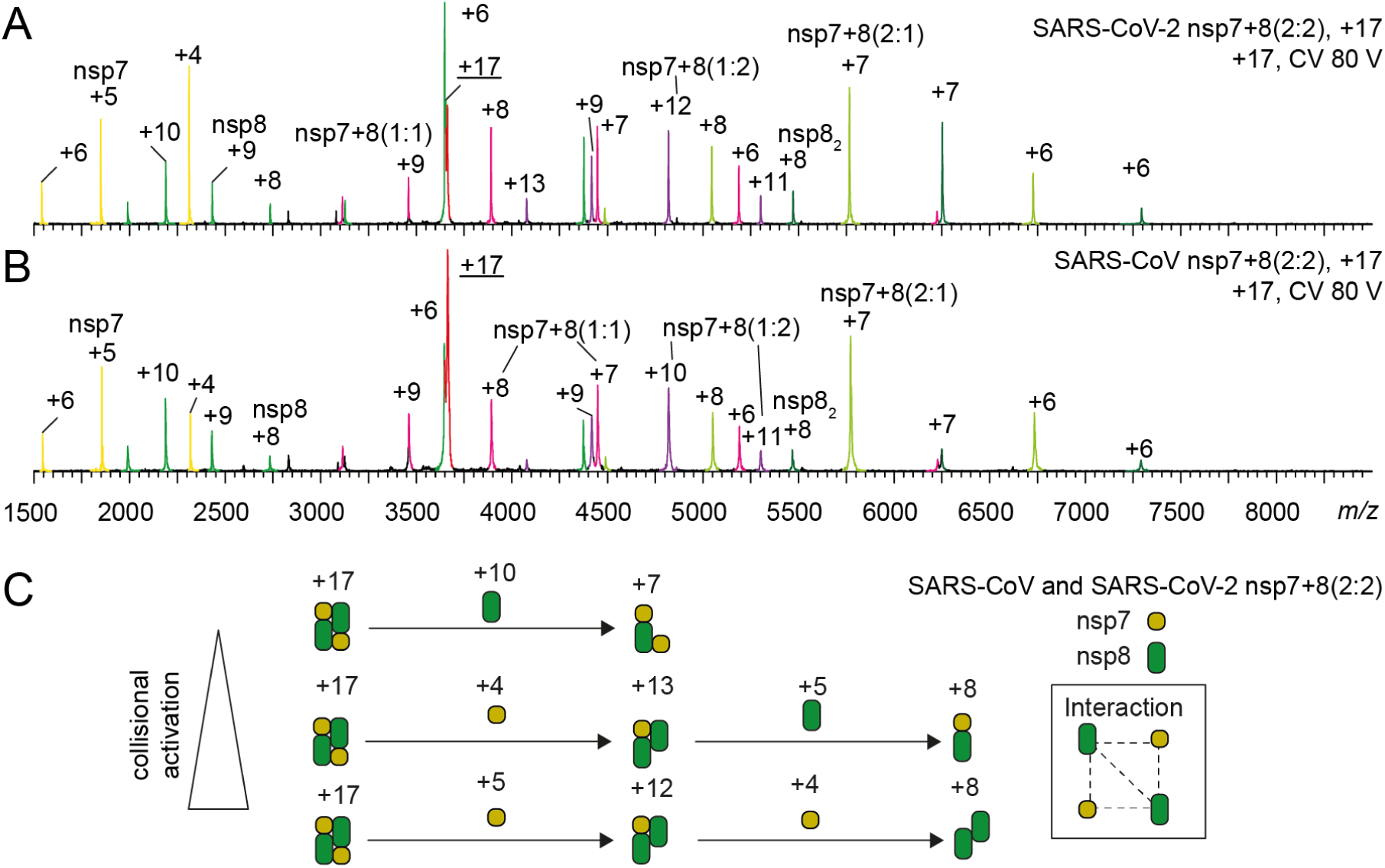
Topological reconstruction from CID-MS/MS of SARS-CoV and SARS-CoV-2 nsp7+8 heterotetramers. Product ion spectra of (**A**) SARS-CoV-2 and (**B**) SARS-CoV 17+ nsp7+8 (2:2) heterotetramers are shown. High charge state precursor allows for efficient dissociation already at relatively low activation (CV 80 V). Product ion species can be clearly assigned from the overview spectra. (**C**) Schematic pathway of dissociation allows topological reconstruction of the SARS-CoV-2 and SARS-CoV heterotetramers.

**Figure S 11:**
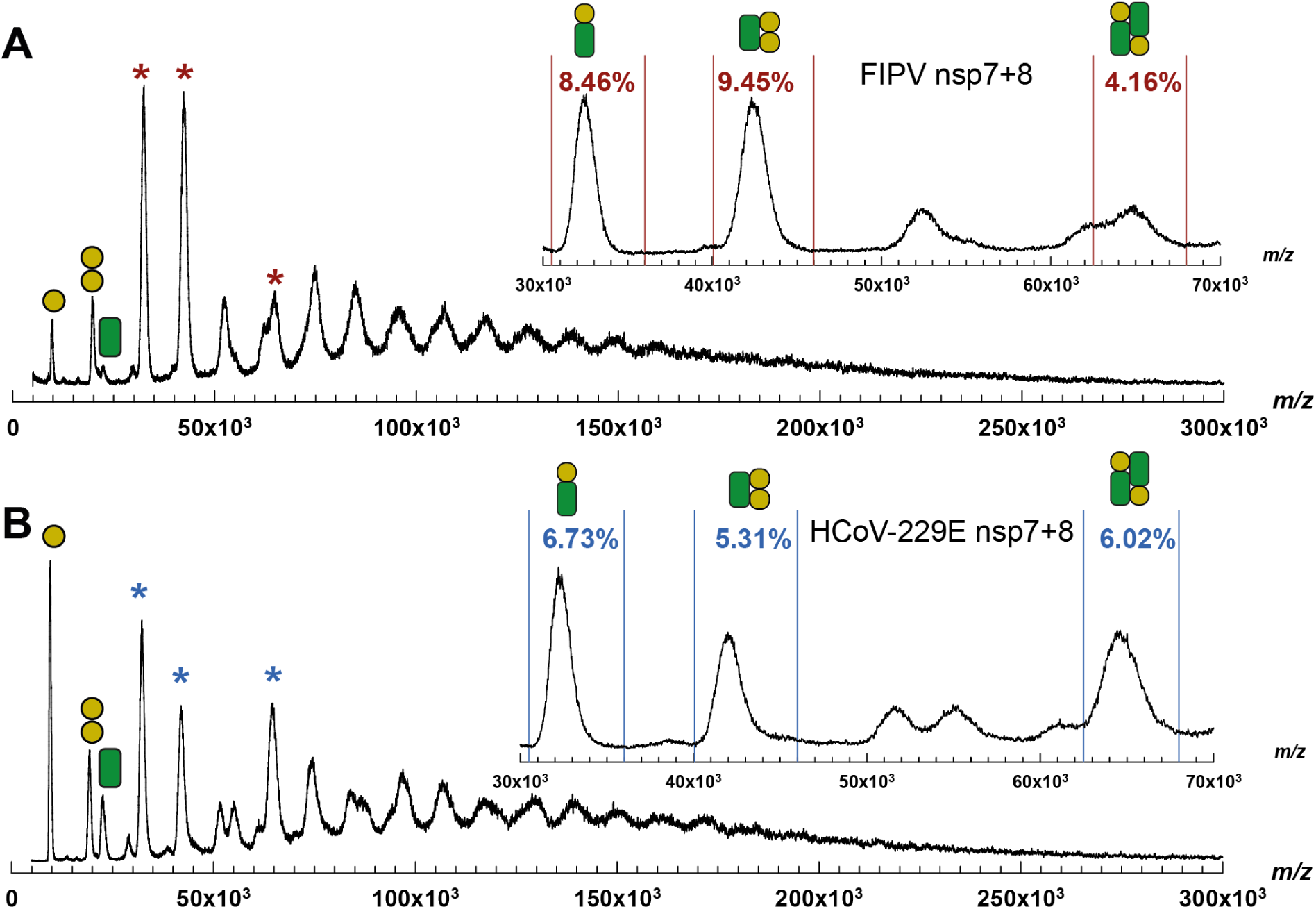
MALDI-MS of nsp7+8 complexes from FIPV and HCoV-229E stabilized with cross-linker. (**A**) MALDI mass spectrum of FIPV nsp7+8 cross-linked with 0.15 % glutaraldehyde for 25 min at 4 °C. Inset shows the most abundant nsp7+8 complexes: (1:1) heterodimer (32.4 kDa) and (2:1) heterotrimer (42.5 kDa). Abundance is determined from relative peak areas as indicated (red). (**B**) MALDI mass spectrum of HCoV-229E nsp7+8 crosslinked with 0.15 % glutaraldehyde for 25 min at 4 °C. Inset shows most abundant nsp7+8 complexes: (1:1) heterodimer (32.2 kDa) and (2:2) heterotetramer (64.6 kDa). Abundance is determined from relative peak areas as indicated (blue). Symbols depict stoichiometry of mass species of nsp7 (yellow) and nsp8 (green) with highest signal strength. Masses are higher in crosslinked samples due to the additional glutaraldehyde molecules. Mass spectra were not calibrated. Each spectrum shown was generated from three MALDI spots. Most signals, except for the HCoV-229E heterotetramer, above 50,000 *m*/*z* are low abundant and likely due to over-crosslinking.

**Figure S 12:**
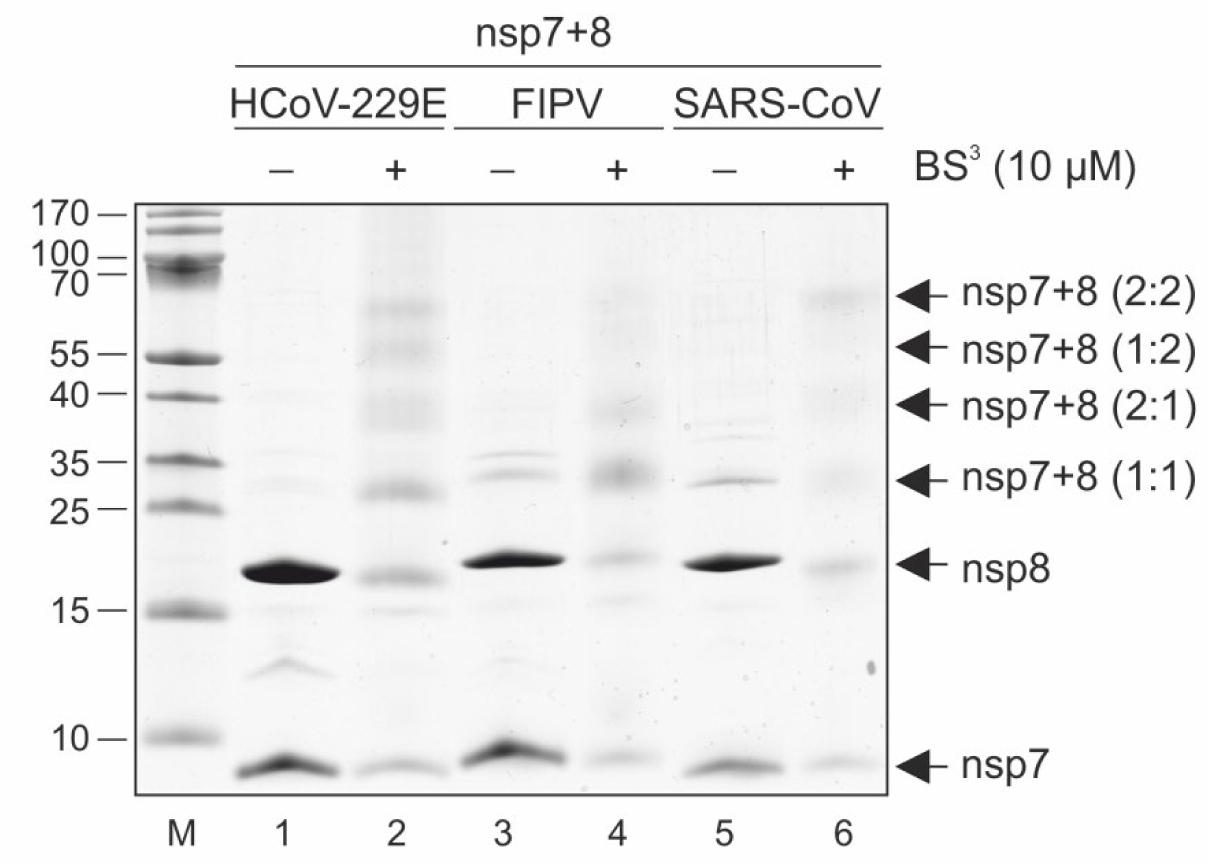
SDS-PAGE analysis of chemically cross-linked HCoV-229E, FIPV, and SARS-CoV nsp7+8 complexes. The 5 μg protein of nsp7+8 complexes were crosslinked with 10 μM BS^3^ (ThermoFisher) in reaction buffer (20 mM HEPES-KOH, pH 8.0, 30 mM KCl, and 2 mM β-mercaptoethanol). Crosslinking was carried out at 37 °C for 30 min and quenched with 50 mM AmAc for another 30 min at 37 °C. After terminating the crosslinking reaction, the samples were mixed with an excess of Laemmli sample buffer (50 mM Tris-HCl, pH 6.8, 2.5 % (*w/v*) SDS, 10 % (*v*/*v*) glycerol, and 0.01 % (*w/v*) bromophenol blue) and analyzed on 12 % SDS-PAGE. Lanes 1, 3, 5 - nsp7+8 complexes not treated with BS^3^ (-) and lanes 2, 4, 6 – nsp7+8 complexes treated with BS^3^. Lane M, marker proteins; molecular masses in kD are indicated to the left. Black arrows on the right indicate the different oligomeric states of the nsp7+8 complexes obtained by crosslinking.

**Figure S 13:**
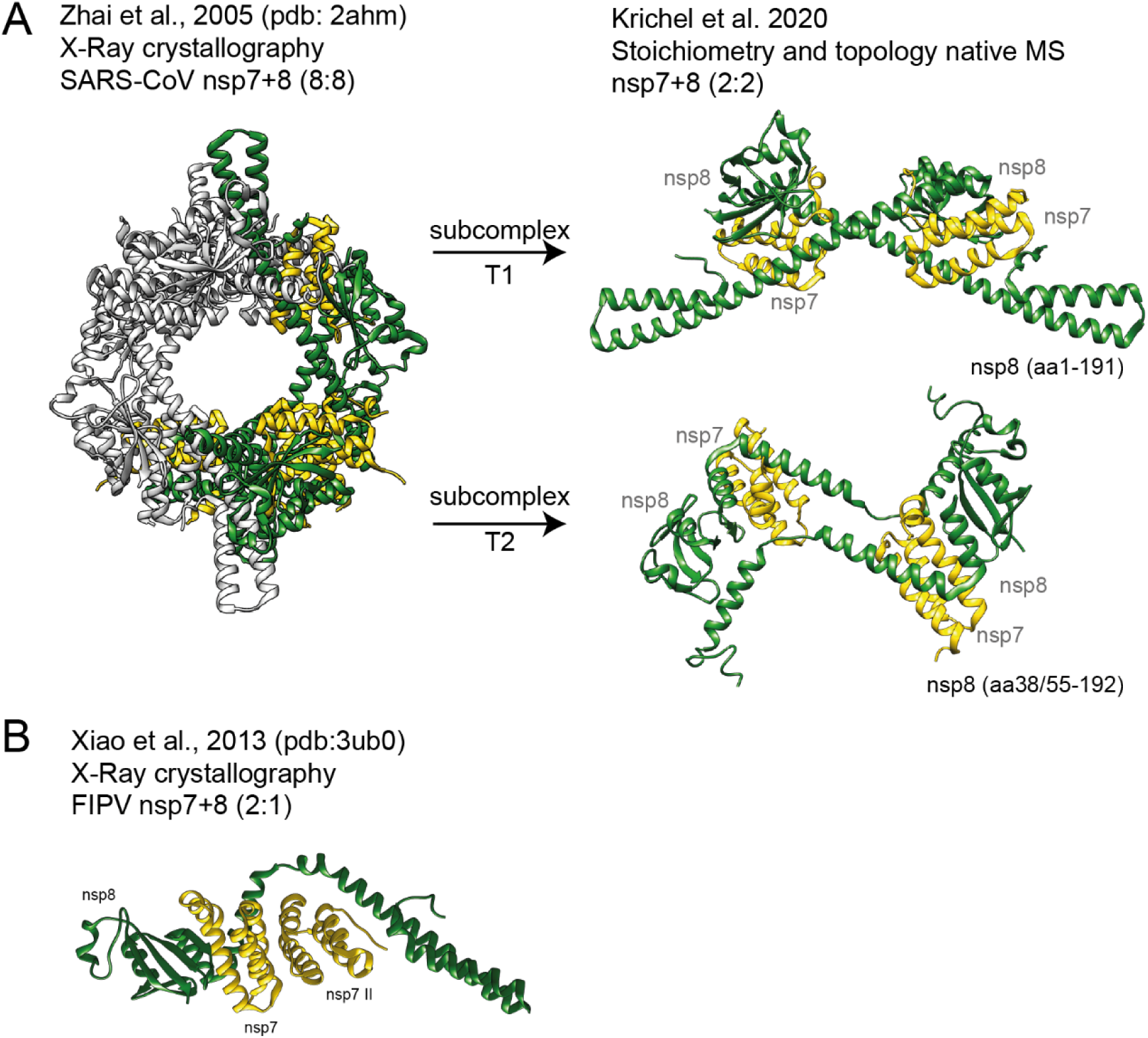
Candidate structures in agreement with the observed stoichiometries and topologies observed. (**A**) For the full-length heterotetramer, an isolated structure does not exist. However from the larger SARS-CoV nsp7+8 hexadecamer [20] (pdb 2ahm), two conformer subcomplexes of nsp7+8 (2:2), T1 and T2, can be extracted. Both conformers constitute a head-to-tail interaction of two heterodimers by an nsp8:nsp8 interface. Notably, nsp8 in T1 is more extended, containing an almost full-length amino acid sequence (2-193), while in T2 the nsp8 *N*-terminal 35 to 55 residues are unresolved. (**B**) For the trimeric complexes, the only deposited structure is FIPV nsp7+8 (2:1) trimer [21] (pdb 3ub0), which agrees well with our experimental topology.

**Figure S 14:**
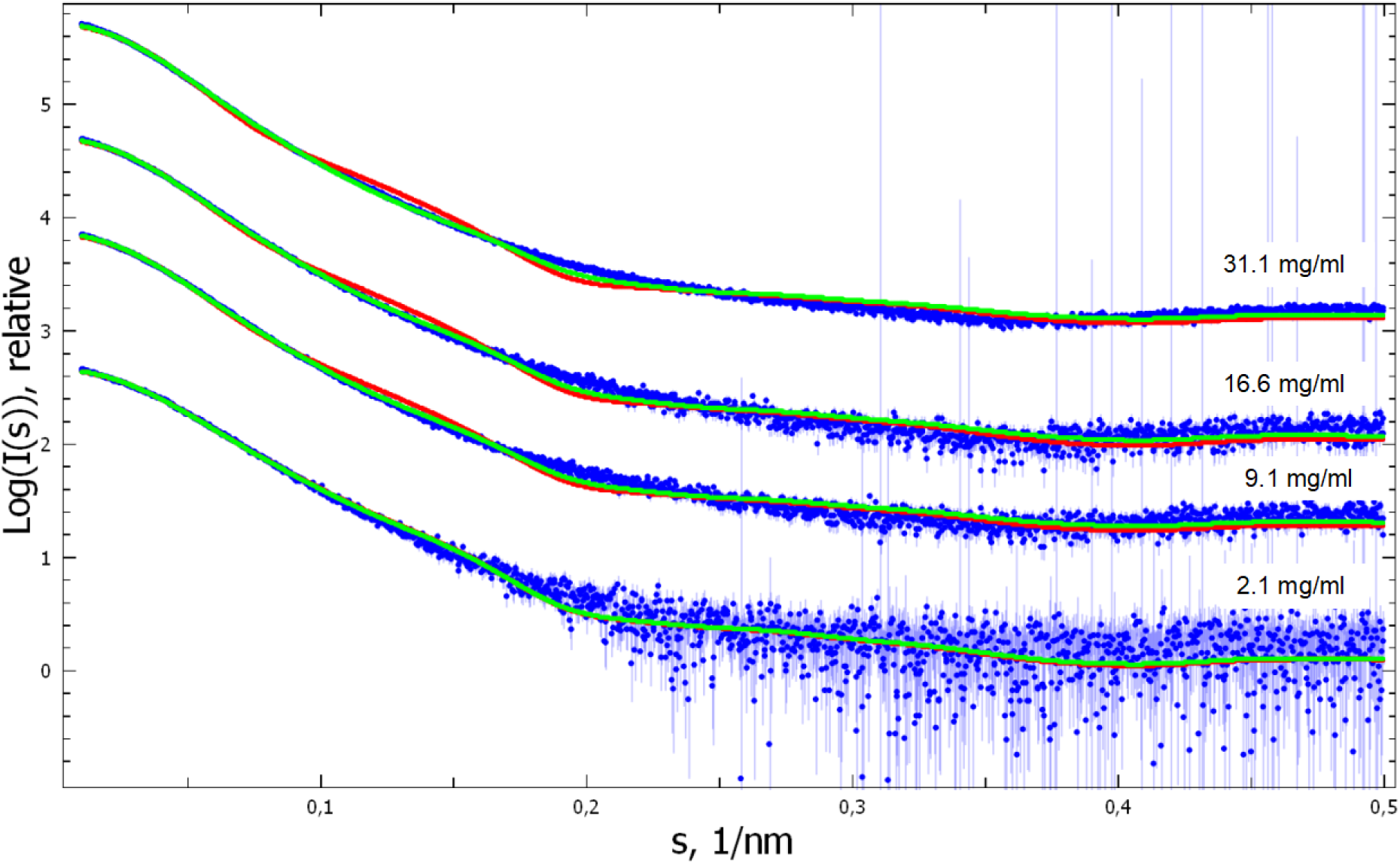
Fits to all SAXS data. Experimental data (blue with experimental errors) for SARS-CoV-2 nsp7+8 complexes fitted with a mixture of T1 and hexadecamer (red), or T1 and dimer of T1 (green).

**Figure S 15:**
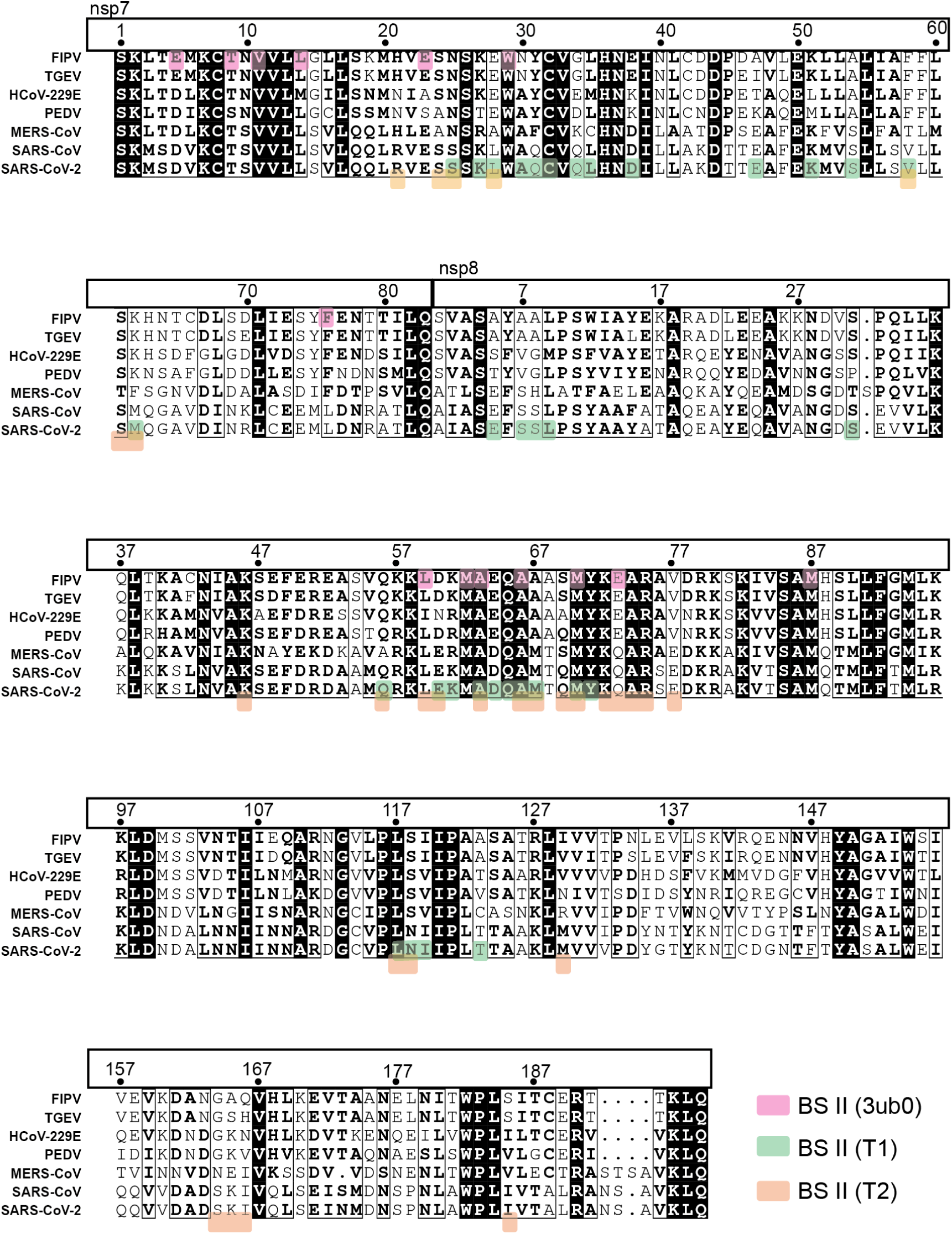
Sequence alignment of nsp7 and nsp8 from seven CoV species. The multiple sequence alignment of nsp7-8 sequences of the seven tested CoVs is genrated with Clustal Omega [43] and converted by ESPript [44] – http://espript.ibcp.fr using the amino acid sequences without C-terminal linkers and His_6_ as input (Table S 6). Highlighted are conserved (black) and semi-conserved (bold) sequences. Further highlighted are molecular contacts at binding site (BS) II in the SARS-CoV nsp7+8 (2:2) heterotetramer candidate structures T1 (green) and T2 (orange) (subcomplexes of pdb 2ahm) as well as in the FIPV nsp7+8 (2:2) heterotrimer (pink, pdb 3ub0). Contacts (VDW radius −0.4 Å) was analyzed with ChimeraX [45].

**Figure S 16:**
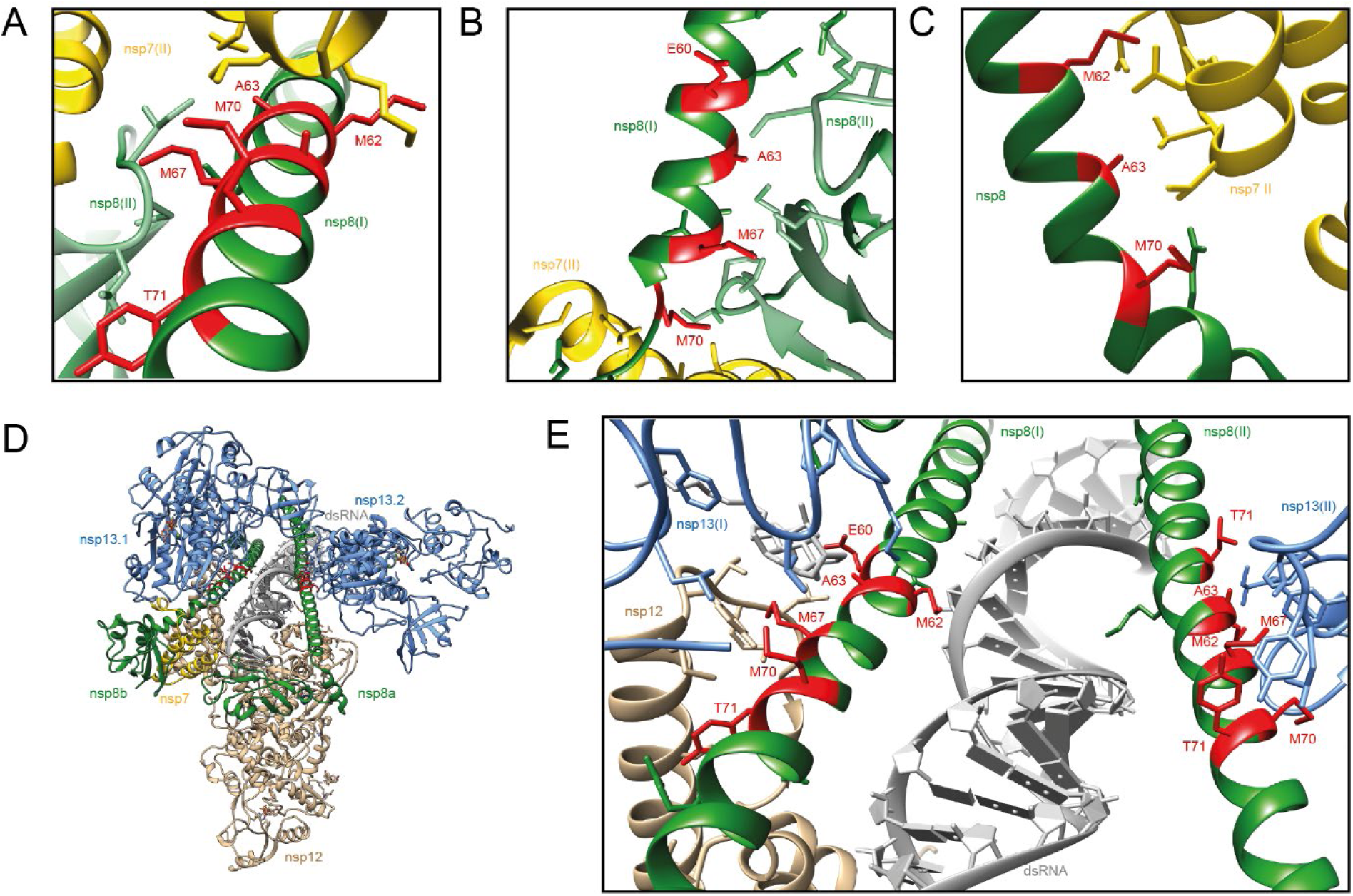
Conserved stretch of nsp8 has various binding contexts. Candidate complexes involving similar conserved residues (red) in the nsp8 BS II are shown here for (**A**) the SARS-CoV nsp7+8 (2:2) heterotetramer candidate structures T1 and (**B**) T2 (subcomplexes of pdb 2ahm) as well as (**C**) the FIPV nsp7+8 (2:2) heterotrimer (pdb 3ub0). (**D**) Cryo electron microscopy structure of the nsp7+8+12+13 (1:2:1:1) polymerase complex (pdb 6xez) [29]. The two subunits of nsp8 (green), bind as a monomer nsp8a and as a nsp7+nsp8b heterodimer to the nsp12 RdRp. From there, nsp8a and nsp8b extend and interact as a facet with the RNA duplex and also individually with one subunit of the nsp13 helicase. The molecular contacts between nsp8 and nsp12/13 are mediated by exactly the same amino acids in the nsp8 BS II that are involved in nsp7+8 complex formation. (**E**) Zoom into the interacting region at nsp8a and nsp8b BS II and its amino acids in contact (VDW radius −0.4 Å) with the nsp12 thumb domain (brown), nsp13.1 and nsp13.2 (blue). All side chains of residues involved in contacts are displayed. Molecular graphics and analyses performed with UCSF ChimeraX, developed by the Resource for Biocomputing, Visualization, and Informatics at the University of California, San Francisco, with support from National Institutes of Health R01-GM129325 and the Office of Cyber Infrastructure and Computational Biology, National Institute of Allergy and Infectious Diseases [45].

## References

1. Cui, J., F. Li, and Z.-L. Shi, Origin and evolution of pathogenic coronaviruses. Nature Reviews Microbiology, 2019. 17(3): p. 181–192.

2. Zaki, A.M., et al., Isolation of a novel coronavirus from a man with pneumonia in Saudi Arabia. New England Journal of Medicine, 2012. 367(19): p. 1814–1820.

3. Drosten, C., et al., Identification of a novel coronavirus in patients with severe acute respiratory syndrome. New England Journal of Medicine, 2003. 348(20): p. 1967–1976.

4. Novel, C.P.E.R.E., The epidemiological characteristics of an outbreak of 2019 novel coronavirus diseases (COVID-19) in China. Zhonghua liu xing bing xue za zhi= Zhonghua liuxingbingxue zazhi, 2020. 41(2): p. 145.

5. Gorbalenya, A.E., Severe acute respiratory syndrome-related coronavirus–The species and its viruses, a statement of the Coronavirus Study Group. bioRxiv, 2020.

6. Dong, E., H. Du, and L. Gardner, An interactive web-based dashboard to track COVID-19 in real time. The Lancet infectious diseases, 2020. 20(5): p. 533–534.

7. Pedersen, N.C., An update on feline infectious peritonitis: virology and immunopathogenesis. The Veterinary Journal, 2014. 201(2): p. 123–132.

8. Vlasova, A., et al., Porcine coronaviruses, in Emerging and Transboundary Animal Viruses. 2020, Springer. p. 79–110.

9. Snijder, E.J., E. Decroly, and J. Ziebuhr, The Nonstructural Proteins Directing Coronavirus RNA Synthesis and Processing. Adv Virus Res, 2016. 96: p. 59–126.

10. van Hemert, M.J., et al., SARS-coronavirus replication/transcription complexes are membrane-protected and need a host factor for activity in vitro. PLoS Pathog, 2008. 4(5): p. e1000054.

11. Shannon, A., et al., Favipiravir strikes the SARS-CoV-2 at its Achilles heel, the RNA polymerase. bioRxiv, 2020.

12. Subissi, L., et al., One severe acute respiratory syndrome coronavirus protein complex integrates processive RNA polymerase and exonuclease activities. Proceedings of the National Academy of Sciences of the United States of America, 2014. 111(37): p. E3900–E3909.

13. Kirchdoerfer, R.N. and A.B. Ward, Structure of the SARS-CoV nsp12 polymerase bound to nsp7 and nsp8 co-factors. Nat Commun, 2019. 10(1): p. 2342.

14. Gao, Y., et al., Structure of the RNA-dependent RNA polymerase from COVID-19 virus. Science, 2020. 368(6492): p. 779–782.

15. Peng, Q., et al., Structural and Biochemical Characterization of the nsp12-nsp7-nsp8 Core Polymerase Complex from SARS-CoV-2. Cell Reports, 2020. 31(11): p. 107774.

16. Hillen, H.S., et al., Structure of replicating SARS-CoV-2 polymerase. bioRxiv, 2020.

17. Ferron, F., et al., Structural and molecular basis of mismatch correction and ribavirin excision from coronavirus RNA. Proceedings of the National Academy of Sciences, 2018. 115(2): p. E162–E171.

18. Gordon, C.J., et al., The antiviral compound remdesivir potently inhibits RNA-dependent RNA polymerase from Middle East respiratory syndrome coronavirus. J Biol Chem, 2020. 295(15): p. 4773–4779.

19. Krichel, B., et al., Processing of the SARS-CoV pp1a/ab nsp7-10 region. Biochem J, 2020.

20. Zhai, Y., et al., Insights into SARS-CoV transcription and replication from the structure of the nsp7-nsp8 hexadecamer. Nat Struct Mol Biol, 2005. 12(11): p. 980–6.

21. Xiao, Y., et al., Nonstructural proteins 7 and 8 of feline coronavirus form a 2:1 heterotrimer that exhibits primer-independent RNA polymerase activity. J Virol, 2012. 86(8): p. 4444–54.

22. Li, S., et al., New nsp8 isoform suggests mechanism for tuning viral RNA synthesis. Protein Cell, 2010. 1(2): p. 198–204.

23. Konkolova, E., et al., Structural analysis of the putative SARS-CoV-2 primase complex. Journal of Structural Biology, 2020. 211(2): p. 107548.

24. Dülfer, J., et al., Structural mass spectrometry goes viral. Advances in virus research, 2019. 105: p. 189–238.

25. Erba, E., L. Signor, and C. Petosa, Exploring the structure and dynamics of macromolecular complexes by native mass spectrometry. Journal of Proteomics, 2020: p. 103799.

26. Wang, G., A.J. Johnson, and I.A. Kaltashov, Evaluation of electrospray ionization mass spectrometry as a tool for characterization of small soluble protein aggregates. Analytical chemistry, 2012. 84(3): p. 1718–1724.

27. Ziebuhr, J., The coronavirus replicase, in Coronavirus replication and reverse genetics. 2005, Springer. p. 57–94.

28. Hegyi, A. and J. Ziebuhr, Conservation of substrate specificities among coronavirus main proteases. J Gen Virol, 2002. 83(Pt 3): p. 595–9.

29. Chen, J., et al., Structural basis for helicase-polymerase coupling in the SARS-CoV-2 replication-transcription complex. Cell, 2020.

30. Akabayov, B., et al., Impact of macromolecular crowding on DNA replication. Nature communications, 2013. 4(1): p. 1–10.

31. Von Brunn, A., et al., Analysis of intraviral protein-protein interactions of the SARS coronavirus ORFeome. PloS one, 2007. 2(5): p. e459.

32. Falke, S., (Doctoral Thesis) Coronaviral Polyprotein Nsp7-10: Proteolytic Processing and Dynamic Interactions within the Transcriptase/Replicase Complex. 2014, Staats-und Universitätsbibliothek Hamburg.

33. Xue, X., et al., Production of authentic SARS-CoV M pro with enhanced activity: application as a novel tag-cleavage endopeptidase for protein overproduction. Journal of molecular biology, 2007. 366(3): p. 965–975.

34. van den Heuvel, R.H., et al., Improving the performance of a quadrupole time-of-flight instrument for macromolecular mass spectrometry. Anal Chem, 2006. 78(21): p. 7473–83.

35. Marty, M.T., et al., Bayesian deconvolution of mass and ion mobility spectra: from binary interactions to polydisperse ensembles. Analytical chemistry, 2015. 87(8): p. 4370–4376.

36. Strohalm, M., et al., mMass data miner: an open source alternative for mass spectrometric data analysis. Rapid Communications in Mass Spectrometry: An International Journal Devoted to the Rapid Dissemination of Up-to-the-Minute Research in Mass Spectrometry, 2008. 22(6): p. 905–908.

37. Franke, D., C.M. Jeffries, and D.I. Svergun, Correlation Map, a goodness-of-fit test for one-dimensional X-ray scattering spectra. Nature methods, 2015. 12(5): p. 419.

38. Franke, D., A.G. Kikhney, and D.I. Svergun, Automated acquisition and analysis of small angle X-ray scattering data. Nuclear Instruments and Methods in Physics Research Section A: Accelerators, Spectrometers, Detectors and Associated Equipment, 2012. 689: p. 52–59.

39. Hajizadeh, N.R., et al., Consensus Bayesian assessment of protein molecular mass from solution X-ray scattering data. Scientific reports, 2018. 8(1): p. 1–13.

40. Svergun, D., C. Barberato, and M.H. Koch, CRYSOL–a program to evaluate X-ray solution scattering of biological macromolecules from atomic coordinates. Journal of applied crystallography, 1995. 28(6): p. 768–773.

41. Konarev, P.V., et al., PRIMUS: a Windows PC-based system for small-angle scattering data analysis. Journal of applied crystallography, 2003. 36(5): p. 1277–1282.

42. Petoukhov, M.V., et al., New developments in the ATSAS program package for small-angle scattering data analysis. Journal of applied crystallography, 2012. 45(2): p. 342–350.

43. Sievers, F., et al., Fast, scalable generation of high-quality protein multiple sequence alignments using Clustal Omega. Molecular systems biology, 2011. 7(1): p. 539.

44. Robert, X. and P. Gouet, Deciphering key features in protein structures with the new ENDscript server. Nucleic acids research, 2014. 42(W1): p. W320–W324.

45. Goddard, T.D., et al., UCSF ChimeraX: Meeting modern challenges in visualization and analysis. Protein Science, 2018. 27(1): p. 14–25.

